# Controlled by disorder: phosphorylation modulates SRSF1 domain availability for spliceosome maturation

**DOI:** 10.1101/2024.12.14.628517

**Authors:** Talia Fargason, Erin Powell, Naiduwadura Ivon Upekala De Silva, Trenton Paul, Peter Prevelige, Jun Zhang

## Abstract

Serine/arginine-rich splicing factor 1 (SRSF1) is key in the mRNA lifecycle including transcription, splicing, nonsense-mediated decay, and nuclear export. Consequently, its dysfunction is linked to cancers, viral evasion, and developmental disorders. The functionality of SRSF1 relies on its interactions with other proteins and RNA molecules. These processes are regulated by phosphorylation of its unstructured arginine/serine-rich tail (RS). Here, we characterize how phosphorylation affects SRSF1’s protein and RNA interaction and phase separation. Using NMR paramagnetic relaxation enhancement and chemical shift perturbation, we find that when unphosphorylated, SRSF1’s RS interacts with its first RNA-recognition motif (RRM1). Phosphorylation of RS decreases its interactions with RRM1 and increases its interactions with the RNA-binding site. This change in SRSF1’s intramolecular interactions increases the availability of protein-interacting sites on RRM1 and weakens RNA binding of SRSF1. Phosphorylation alters the phase separation of SRSF1 by diminishing the role of arginine in intermolecular interactions. These findings provide an unprecedented view of how SRSF1 influences the early-stage spliceosome assembly.

**SUMMARY:** Phosphorylation of SRSF1 is pivotal in pre-mRNA processing and is dysregulated in various pathologies. Modeling of SRSF1 based on NMR restraints reveals phosphorylation alters the accessibility of protein-protein and protein-RNA interaction sites on SRSF1’s RRM1 domain, altering its binding preferences

## INTRODUCTION

In eukaryotic genes, coding sequences (exons) are interrupted by non-coding sequences (introns), making RNA splicing an essential process to remove introns and join exons together. RNA splicing is performed by the spliceosome — a dynamic, multi-component complex consisting of small nuclear ribonucleoproteins (U1, U2, U4, U5, and U6) (Fig. 1A). This process is regulated by and relies on splicing factors. Ser/Arg-rich proteins (SR proteins) are a key family of twelve splicing factors characterized by one to two RNA-recognition motifs (RRMs) and a C-terminal Arg/Ser-rich tail (RS). As the founding member of the SR family, SRSF1 plays crucial roles in splicing, maintaining genome stability,^1^ promoting mRNA transcription,^2, 3^ transportation,^4^ and translation,^5, 6^ nonsense-mediated mRNA decay,^7^ immune response,^8^ and interactions with long noncoding RNA.^9–13^ As a results, their dysregulation is associated with various types of cancer,^14^ viral evasion,^15,16^ neurodevelopmental conditions,^17,18^ and neurodegenerative disorders. ^19^

**Figure 1:**
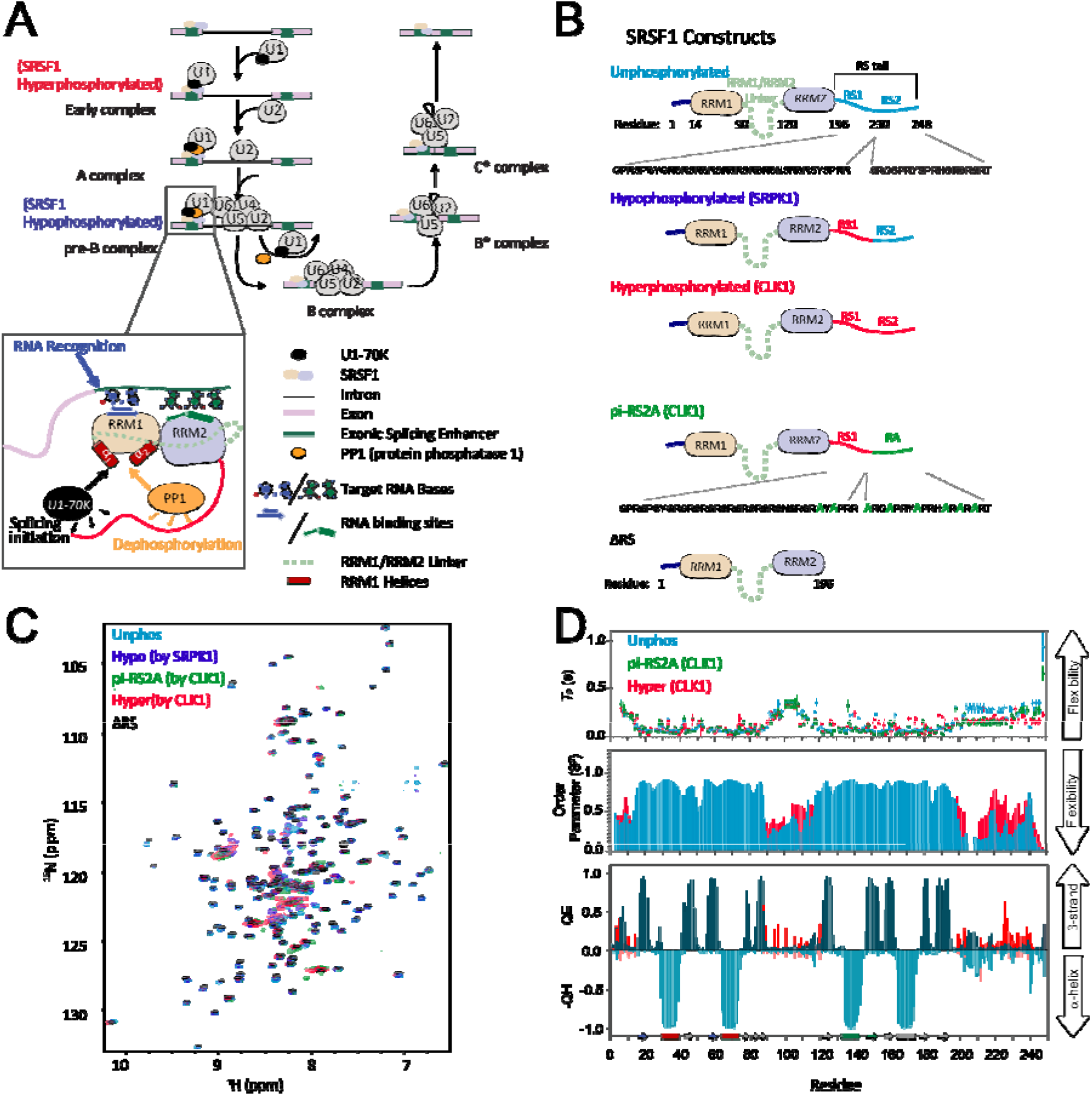
SRSF1’s linker and tail are unstructured but can form transient structures. (A) SRSF1’s role in the splicing process. Dephosphorylation of SRSF1 between the early and the B complex is considered necessary for spliceosome activation. Crucial ligand binding sites are expanded in the inset; on the RRM1 domain, RNA-binding occurs on the beta sheet regions while protein-protein interactions occur on the helices on the opposite side of the RRM1 domain. (B) SRSF1 constructs used in this paper. SRSF1 was expressed and purified in its three major phosphorylation states. Artificial pi-RS2A and ΔRS constructs were also used to gain further insight into the key differences between the hypo- and hyper-phosphorylated states as well as the effect of adding an RS domain in any state, respectively. (C) HSQC NMR spectral overlay of SRSF1 constructs. (D) SRSF1 is a partially disordered protein. ^15^N amide transverse NMR relaxation time (*T*_2_), chemical shift-derived order parameters (*S*^2^) and secondary structure propensities, QE for β-strand, and -QH for α-helix.

SRSF1, like other SR proteins, promotes the spliceosome assembly around the 5’ splicing site. SRSF1 fulfills this role by serving two functions: binding to RNA elements known as exonic splicing enhancers and mediating protein-protein interactions. SRSF1 recognizes exonic splicing enhancers through its RRM1 and RRM2 domains (Fig. 1A, 1B). RRM1 preferentially binds to C-containing motifs, while RRM2 primarily recognizes guanine-rich motifs, such as GGA.^20^ RNA binding to RRM1 occurs on the β1 and β3 strands of the domain,^20^ while on RRM2, it happens at the edge of the α1 helix and β2 strand. SRSF1 directly interacts with U1-component protein U1-70K, which recruits the U1 complex to the splicing site (Fig. 1A).^21^ The SRSF1/U1-70K interaction is mediated mainly by RS of SRSF1 and assisted by RRM1 through the two RRM1 helices (α1, α2) opposite the RNA-binding site.^22,23, 24^ Notably, these RRM1 helices are also responsible for inter-molecular interaction with protein phosphatase 1 (PP1) and intra-molecular interaction with the RS tail.^22,23 25^

SRSF1’s role in splicing, as well as its cellular localization, are regulated by phosphorylation of its RS tail. In the cytoplasm, SRSF1 is phosphorylated by SR-Protein Kinase 1 (SRPK1), which phosphorylates 8-12 consecutive serine residues in the RS tail, generating the hypophosphorylation form.^26^ In this state, SRSF1 binds to nuclear importers to enter the nucleus,^27^ where it undergoes further phosphorylation by SRPK1 and CLK1, resulting in a hyperphosphorylated form (18-22 phosphorylated serines).^26,28^ Phosphorylation also regulates SRSF1’s role in the spliceosome assembly. Hyper-phosphorylation of SRSF1 is critical for its interaction with U1-70K.^22,23, 24^ However, partial dephosphorylation of SRSF1 is needed to activate the spliceosome (Fig. 1A).^22, 29^ Therefore, the balance of phosphorylation/dephosphorylation orchestrated by SRPK1, CLK1, and PP1 is critical to maintaining cellular homeostasis.^23^ Hyperactive kinases are linked to cancers,^30,31,32^ but inhibition of the same kinases under the wrong circumstances is also linked to chemoresistance.^33,34,35,36^

Phosphorylation also governs SRSF1’s cellular localization to nuclear speckles, a type of membraneless organelle driven by liquid-liquid phase separation (LLPS). Over expression of CLK1 results in release of SRSF1 from the nuclear speckles.^37^ Our previous study showed that LLPS of SRSF1 relies on its RS tail.^25^ These results together suggest that phosphorylation of RS regulates phase separation behavior of SRSF1.

This study aimed to elucidate the structural changes that accompany SRSF1’s transition from unphosphorylated to phosphorylated states. Paramagnetic relaxation enhancement (PRE) NMR revealed that in its unphosphorylated state, the RS tail interacts with the α1 and α2 helices of RRM1 that are responsible for protein-protein interaction. As SRSF1 becomes phosphorylated, the RS tail engages less with these sites and more with other regions of RRM1, including the RNA-binding site. The result of this change in intramolecular interactions is that responsible for U1-70K and PP1 interaction, are autoinhibited by the unphosphorylated RS tail and become accessible upon phosphorylation, facilitating U1-70K binding. Our NMR and molecular dynamics simulations suggest that the Arg residues in the RS tail form salt bridges with adjacent serines. In combination with our in vitro phase separation assays, our work also provides a mechanism by which phosphorylation regulates SRSF1’s phase separation propensity. These insights into how phosphorylation regulates SRSF1 structure, RNA binding, U1-70K interaction, and phase separation shed light on splicing regulation and potential therapeutic interventions.

## MATERIALS AND METHODS

### Molecular cloning and Protein expression

The DNA encoding human SRSF1 (residues 1-248) was sub-cloned into pSMT3 using BamH I and Hind III. Human Clk1 and SRPK1 were sub-cloned into pETDuet-1 MCS1 (Nco I/Sac I) and MCS2 (EcoR V/Kpn I), respectively. SRSF1 constructs SRSF1^1–109^, SRSF1^110–196^, SRSF1^1–196^, SRSF1^197–248^ and its mutants C16S/C148S/N220C, C16S/C148S/T248C, RS2A (phosphorylatable serines on residues 225-248 mutated to alanine) were prepared using mutagenesis PCR.

All proteins were expressed by BL21-CodonPlus (DE3) cells in LB media or minimal media supplemented with proper isotopes. To produce phosphorylated SRSF1, BL21-CodonPlus cells were co-transformed by pSMT3/SRSF1 and pETDuet-1/Clk1/SRPK1. Cells were cultured at 37 °C to reach an OD_600_ of 0.6, when 0.5 mM IPTG was added to induce protein expression. Cells were further cultured 16-20 hours at 22 °C. The cells were harvested by centrifugation (4000 RCF, 15 min) and stored in −80 °C.

### Deuterated Cell growth for Purification of ^2^H ^13^C ^1 5^N SRSF1 (residues 1-248)

Two liters of deuterated M9 media were prepared with ^15^N Ammonium Sulfate and ^13^C glucose using pure D_2_O as a solvent. Cells were slowly introduced to deuterated M9 medium by increasing the ratio of D_2_O to H_2_O in increments. A 3 mL LB/antibiotic mixture was inoculated with a colony of cells and allowed to shake at 200 RPM, 37 °C for two hours. Next, 0.75 mL of the resulting LB growth was transferred into a container of 2.25 mL deuterated M9 medium (resulting in a 75% D_2_O/25% H_2_O culture) and allowed to grow for 5 hours. Once this time had elapsed, the growth was transferred to a shaker flask with 22 mL deuterated M9 culture (resulting in 85% D_2_O/15% H_2_O) and allowed to grow for 16 hours. After this, 12.5 mL of the resulting culture was evenly distributed between four shakers containing 100 mL deuterated M9 media each and shaken until an OD_600_ of 0.6 was reached. These 100 mL growths were then transferred to flasks containing 400 mL deuterated M9 media and grown at 37 °C until an OD_600_ of 0.8 was reached. At this point, IPTG was added to a concentration of 1 mM, and cells were shaken at 22 °C for 16 hours before harvesting.

### Protein purification

*SRSF1(residues 1-196):* The cell pellet was re-suspended in 20 mM Tris-HCl, pH 7.5, 2 M NaCl, 25 mM imidazole, 0.2 mM TCEP supplemented with 1 mM PMSF, 0.5 mg/mL lysozyme, and 1 tablet of Protease Inhibitor. After three freeze-thaw cycles, the sample was sonicated and centrifuged at 23,710 g for 40 min using a Bruker Avanti JXN26/JA20 centrifuge. The supernatant was loaded onto 5 mL HisPur Nickel-NTA resin and washed with 200 mL 20 mM Tris-HCl, pH 7.5, 2 M NaCl, 25 mM imidazole and 0.2 mM TCEP. The protein was then eluted with 30 mL 20 mM MES pH 6.5, 500 mM imidazole, 500 mM Arg/Glu, 0.2 mM TCEP. The eluted sample was cleaved with 2 µg/mL Ulp1 for 2 hours at 37 °C, diluted three-fold with a buffer A of 20 mM MES pH 6.0, 50 mM arginine/glutamate, 0.1 mM TCEP, and loaded onto a 5-mL HiTrap Heparin column. The sample was eluted over a gradient with a buffer B of 20 mM MES pH 6.0, 50 mM arginine/glutamate, 0.1 mM TCEP, 2 M NaCl, 0.02% NaN_3_. SRSF1^1–196^ were eluted around 50% B. Fractions containing SRSF1^1–196^ were pooled, concentrated, and loaded onto a HiLoad 16/60 Superdex 75 pg size exclusion column equilibrated with 0.2 M Arg/Glu, 0.2 M NaCl, 20 mM Tris-HCl, pH 7.5, 0.1 mM TCEP, 0.02% NaN_3_.

### Unphosphorylated SRSF1(residues 1-248)

Once pelleted, cells were resuspended in a lysis buffer of 20 mM Tris-HCl pH 7.5, 150 mM Arg/Glu, 25 mM imidazole, 2 M NaCl, 1 mM TCEP, 1 mg/mL lysozyme, 1 mM PMSF, and 1 tablet Pierce protease inhibitor. They were then subjected to three freeze-thaw cycles and sonication. After this point, all steps occurred at 22 °C or higher. The lysate was spun at 25 °C, 23,710 g for 30 minutes using a Bruker Avanti JXN26/JA20 centrifuge. The supernatant was applied to a 5 mM nickel column that was then washed with 200 mL high salt buffer (1 M Urea, 25 mM imidazole pH 8.0, 1 mM TCEP, 4.3 M NaCl) followed by 100 mL Loading Buffer (20 mM Tris-HCl pH 7.5, 25 mM imidazole, 1 M NaCl, 1 mM TCEP). The sample was eluted using 30 mL elution buffer (500 mM Arg/Glu pH 6.5, 250 mM imidazole, 1 mM TCEP, 1 Pierce protease inhibitor tablet), supplemented with 1 mM PMSF and 2 µg/mL Ulp1, then incubated at 37 °C for 2 hours. The sample was diluted 3-fold using a Heparin Buffer A of 100 mM Arg/Glu, pH 4.6, 0.02% NaN_3_, 1 mM TCEP, where pH was adjusted as necessary with acetic acid. The sample was then loaded to a 5 mL HiTrap Heparin column and eluted over a gradient with a Buffer B of 1 M Arg/Glu pH 8.5, 1 M NaCl, 0.02% NaN_3_, 1 mM TCEP.

### Hyperphosphorylated and pi-RS2A SRSF1(residues 1-248)

Mutants of SRSF1 1-248 were co-expressed with the phosphokinase CLK1. Once pelleted, cells were resuspended in a lysis buffer of 20 mM Tris-HCl pH 7.5, 100 mM Arg/Glu, 25 mM imidazole, 2 M NaCl, 1 mM TCEP, 1 mg/mL lysozyme, 1 mM PMSF, 1 Pierce Protease inhibitor tablet, and 1 mM Na_3_VO_4_. Cells were then lysed through a combination of three freeze-thaw cycles and sonication. The cell lysate was spun at 4 °C, 23,710 g for 30 minutes using a Bruker Avanti JXN26/JA20 centrifuge. After this point, all steps occurred at 22 °C or higher. The supernatant was applied to a 5 mL nickel column that was then washed with 200 mL high salt buffer (1 M guanidinium, 25 mM imidazole pH 8.0, 1 mM TCEP, 4.3 M NaCl) followed by 20 mL low salt buffer (20 mM Tris-HCl pH 8.0, 25 mM imidazole, 1 mM TCEP). The protein was then eluted using 30 mL elution buffer (500 mM Arg/Glu pH 8.5, 500 mM imidazole, 1 mM NaVO_4_, 1 mM TCEP, 1 Pierce Protease inhibitor tablet. The eluate was supplemented with 1 mM PMSF, 1 mM EDTA, and 2 µg/mL Ulp1, and incubated at 37 °C for 2 hours. After cleavage the sample was diluted 3-fold using a Buffer A of 50 mM Arg/Glu pH 8.5, 1 mM EDTA, 0.02% NaN_3_, 1 mM TCEP and loaded to a 5 mL HiTrap Q column. A gradient elution was performed with a Buffer B of 100 mM Arg/Glu pH 8.5, 2M NaCl, 1 mM EDTA, 0.02% NaN_3_, 1 mM TCEP.

### Hypophosphorylated SRSF1(residues 1-248)

Hypophosphorylation of SRSF1 was performed *in vitro* by incubation of 100 µM SRSF1 constructs with 5 µM SRPK1 at 22 °C for 12-16 hours in 20 mM HEPES pH 7.5, 500 mM Arg/Glu, 0.2 mM TCEP, 2 mM ATP, 5 mM MgCl_2_.

### SRSF1 RRM1(residues 14-90) and RRM1 + ½ linker (residues 1-109)

The cell pellet was resuspended in a lysis buffer containing 25 mM HEPES, pH 8.5, 1M NaCl, 25 mM imidazole, 1 mM TCEP, 1 mg/mL lysozyme, 1 tablet of Pierce protease inhibitor, 1 mM PMSF, 0.02% NaN_3_. After three freeze–thaw cycles, the sample was sonicated and centrifuged at 23,710 RCF for 40 min using a Beckman Coulter Avanti JXN26/JA20 centrifuge. The supernatant was loaded onto 5 mL of HisPur Nickel-NTA resin and washed with 100 mL of a loading buffer (25 mM HEPES, pH 8.5, 1M NaCl, 25 mM imidazole, 1 mM TCEP, 0.02% NaN_3_), followed by 100 mL of a high salt wash buffer (5M NaCl, 25 mL imidazole, 1 mM TCEP, and 0.02% NaN_3_), and finally an additional 100 mL of loading buffer. On-column cleavage was performed in 25 mL loading buffer supplemented with 2 μg/mL ULP1, 1 mM PMSF, 1 protease inhibitor tablet through inversion for 2 h. Protein was eluted using an elution buffer of 20 mM Tris-HCl pH 8.5, 500 mM imidazole 1 mM TCEP, 0.02% NaN_3_. The protein was concentrated and loaded onto a HiLoad 16/600 Superdex 75 pg size exclusion column equilibrated with 200 mM Arg/Glu, 0.2M NaCl, 20 mM Tris– HCl, pH 7.5, 0.1 mM TCEP, and 0.02% NaN_3_.

### SRSF1 RRM2(residues 121-196) and RRM2 + ½ linker (residues 110-196)

The cell pellet was resuspended in a lysis buffer (25 mM HEPES, pH 8.5, 1M NaCl, 25 mM imidazole, 1 mM TCEP, 1 mg/mL lysozyme, 1 Pierce protease inhibitor tablet per 30 mL, 1 mM PMSF, 0.02% NaN_3_). After three freeze–thaw cycles, the sample was sonicated and centrifuged at 23,710 RCF for 40 min using a Beckman Coulter Avanti JXN26/JA20 centrifuge. The supernatant was loaded onto 5 mL of HisPur Nickel-NTA resin and washed with 100 mL of a loading buffer (25 mM HEPES, pH 8.5, 1M NaCl, 25 mM imidazole, 1 mM TCEP, 0.02% NaN_3_), followed by 100 mL of a high salt wash buffer (5 M NaCl, 25 mL imidazole, 1 mM TCEP, and 0.02% NaN_3_) and finally an additional 100 mL of loading buffer. The protein was eluted using an elution buffer of 20 mM Tris-HCl pH 8.5, 500 mM imidazole 1 mM TCEP, 0.02% NaN_3_. The protein was further purified using a 5-mL Cytiva Fast Flow Q column on a gradient between a buffer A of 50 mM Arg pH 8.5, 1 mM and a buffer B of 50 mM Arg/Glu pH 8.5, 2M NaCl, 1 mM TCEP. As SRSF1 121-196 was not stained well by Coomassie brilliant blue, an HSQC-NMR spectrum was obtained to verify sample identity and purity (>95%).

### Mass Spectrometry

The protein was diluted to 20 µM into a cleavage buffer of 400 mM Arg/Glu pH 8, 1 mM TCEP, 1 mM EDTA, 0.02% NaN_3_. The samples were cleaved by Lys-C with a protease: SRSF1 ratio of 1:20 at 37 °C for either 10 minutes to cleave unstructured regions (Fig. S1B-E), or 12 hours to cleave structured regions (Fig. S1F-G). At the end of the incubation, Lys-C was inactivated by 250 mM HCl. Digested samples were diluted into 0.1% formic acid and separated on a 2.5 × 150 mm A BEH SEC column (Waters) using 0.1% formic acid at a flow rate of 70 μL/min. The elute of the column was connected inline to the ESI source of a Waters Synapt G2-S(i). Data was collected under MassLynx (Waters) in positive ion, resolution mode and the spectra were deisotoped using the MaxEnt3 module within MassLynx and the mass versus intensity plots. For protease cleavage experiments, intensity values are normalized to the intensity of the highest peak.

### NMR assignment experiments

The SRSF1 construct (residues 1-248) was prepared as described above except that the *E. coli* cells were grown in M9 media containing ^15^N, ^13^C, and ^2^H isotopes. The protein (∼ 370 µM) was purified as described above, exchanged into and concentrated in 800 mM Arg/Glu pH 6.5, and diluted into 100 mM ER4 peptide (amino acid sequence ERER), 400 mM RE pH 6.4, 1 mM TCEP, 5% D_2_O 0.02% NaN_3_ for NMR measurements. Triple resonance assignment experiments HNCA, HNCACB, HN(CO)CA, HN(CO)CACB, HNCO, and MUSIC were collected at 37 °C on a Bruker Avance III-HD 850 MHz spectrometer installed with a cryo-probe. The NMR data was processed using NMRPipe^38^, and assignment was performed using NMRViewJ.^39^

### Chemical Shift Perturbation experiments

HSQC spectra were collected, unless otherwise indicated, at 310K in 200 mM Arg/Glu, 20 mM MES, pH 6.5, 5% D_2_O, on a 600 MHz Bruker Avance III-HD 600 MHz magnet. Chemical shift perturbations were calculated according to the equation:

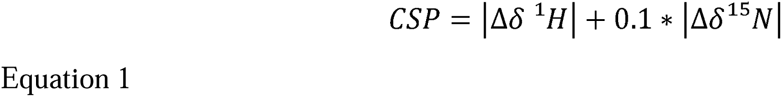

Where Δδ^1^H is the difference in chemical shift values between the two spectra in the proton dimension and Δδ^15^N is the difference in chemical shift values between the two spectra in the nitrogen dimension.

A standard deviation for each peak was calculated according to the equation:

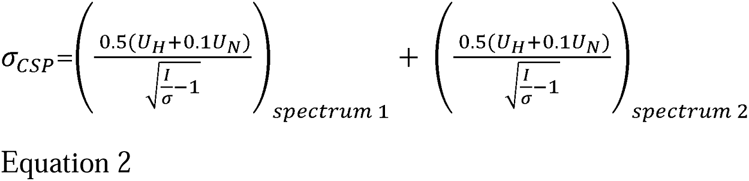

Where *U _H_* and *U _N_* are the linewidths of a given peak in the hydrogen and nitrogen dimensions, respectively, *I* is the intensity of the peak, and *σ* is the standard deviation of the spectrum calculated using NMRViewJ. The line of identity was drawn at the median value of *σ_CSP_* across all peaks.

### Paramagnetic relaxation enhancement

Protein constructs were desalted using a HiPrep 26/10 desalting column into an MTSL labeling buffer of 0.8 M Arg/Glu, 100 mM NaCl, 50 mM Tris-HCl, 1 mM EDTA pH 7.5 and diluted to a concentration of 20 µM. MTSL was added to a concentration of 400 µM. The sample was incubated in the dark at 37 °C for 12 hours, after which the constructs were desalted into a buffer of 0.8 mM Arg/Glu pH 6.3, 1 mM EDTA. The sample was concentrated, after which ER4 peptide stock and D_2_O were added to produce a final NMR buffer of 100 mM ER4, 400 mM Arg/Glu pH 6.3, 0.2 mM EDTA, 5% D_2_O.

Spectra were collected at 37 °C on a Bruker AVANCE 850 MHz NMR spectrometer. Measurements were carried out using a pulse sequence developed by Junji Iwahara.^3^ After collection of paramagnetic spectra, samples were quenched using 10 mM sodium ascorbate, and diamagnetic spectra were obtained. The NMR data was processed using NMRPipe^38^ and analyzed using NMRViewJ.^39^

PRE in the peptide buffer was obtained using a 5-datapoint fitting for the rate of *T_2_* relaxation under paramagnetic and diamagnetic conditions. PRE was calculated as the difference between the rate of decay in the paramagnetic state and that in the diamagnetic state. Error was estimated by adding the error from the exponential fit of the relaxation of spectra obtained in the paramagnetic and diamagnetic states.

### iPRE

The relative PRE was calculated according to the following equations:

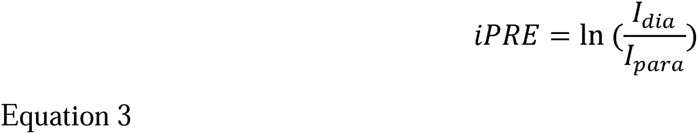

where, *I_dia_* and *I_para_* are the intensities of the individual peak in the diamagnetic and paramagnetic states, respectively.

Error of individual peaks was approximated as:

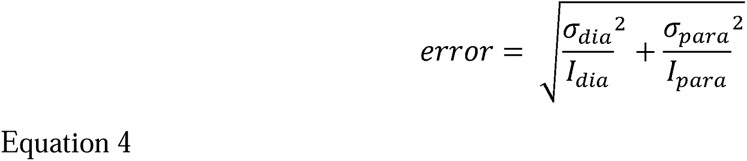

where *σ_dia_* and *σ_para_* are the standard deviations of the diamagnetic and paramagnetic spectra, respectively. For the phosphorylated protein in peptide buffer *σ_dia_* was 0.89, and *σ_para_* was 0.86.

### Molecular Modeling

The Xplor-NIH^40,41^ structures were refined against PRE, CSP, and dihedral angle restraints. Refinement against PRE or iPRE was accomplished by minimizing the Pearson correlation coefficient and Q-value in the SBMF mode.^42^ Ensemble structures with Q values less than 0.4 were combined to create a pool. Dihedral angles were calculated from backbone chemical shifts in the program TALOS^43^ before incorporation into the refinement script through the “convertTalos” helper program. Chemical shift perturbations and bleaching on PRE were incorporated into Xplor-NIH models as artificial NOE constraints.^44^ Bleached residues were assigned as artificial NOE constraints in which the upper limit was set to 12 Å. Distances across an ensemble structure were averaged according to the equation:

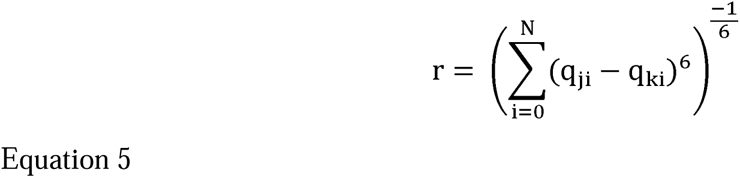

Where N is the total number of ensemble members q_ji_ are the coordinates of atom j in the i^th^ ensemble and q_ki_ are the coordinates of atom k in the i^th^ ensemble. If the calculated average distance was less than or equal to 12 Angstroms, the model was considered to satisfy this constraint. Q-factors of selected structures ranged from 0.148-0.401 (Table S2).

The best-fit ensemble structures (by lowest Q-factor) were selected for refinement in AMBER20 (Fig. S5).^45^ The forcefield ff14IDPSFF^46^ with TIP3P water was used with the phosphorylation forcefield of phosaa14SB where relevant at 310 K. The IDP CMAP parameters were applied only to regions of the protein known to be unstructured. This included residues 1-14, 90-120, and 196-248. Bonding interactions were determined by analysis of endpoint pdb files created through the ambpdb command. Cpptraj was used to identify hydrogen bonding interactions, and stacking interactions were identified through an in-house python script available on our github (https://github.com/taliafargason). Arg-Arg stacking between two arginine residues, pi-pi stacking between two aromatic residues, and cation-pi stacking interactions between arginine and aromatic residues were determined to occur if the sidechains of the residues in question were on planes that were facing each other within an error of 20° and that were within a distance range of 4.5 Å from each other.

### AlphaFold Structures

Structures of PP1γ and U1-70k bound to hyperphosphorylated SRSF1 were constructed using the AlphaFold3 Server.^47^ The best five structures for each interaction were examined in PyMOL and compared to literature CSP and mutagenesis experiments, which identify RRM1 residues involved in binding to both of these ligands.^23,24,22^ Those matching experimental data were selected for further study.

### Autoinhibition

The PyMOL Molecular Graphics System^48^ was used to identify conformers that were autoinhibited against a given binding partner. The ensemble structures presented in Fig. 6 were loaded into a PyMOL session along with an AlphaFold structure of SRSF1 bound to either PP1γ or U1-70K. Each conformer in the ensemble structure was assigned using the RRM1 domain (SRSF1 residues 14-90) as an anchor in the AlphaFold structure. The “select overlapatom” command was used to identify SRSF1 residues in the range of 91-248 that overlapped with a 4 Å surface around the binding partner. (The precise command used was: “select overlapatom, chain B around 4 and not resid 1-90”). A conformer in the ensemble structure was defined as autoinhibited against a binding partner if residues 90-248 were found to overlap with a 4 Å surface around the binding partner. The BAD domains of U1-70k were not included in this analysis. Values in Fig. 7H represent the percentage of conformers in the ensemble structure that meet the definition of autoinhibited.

### Fluorescence polarization (FP) Assays

Fluorescence polarization was obtained using fluorescein-tagged RNA constructs. The protein was combined at the highest indicated concentration with 10 nM RNA probe, and serial dilutions into a binding buffer containing 10 nM of RNA probe were performed to obtain lower concentrations of protein. Binding was determined either in an arginine buffer of 250 mM Arg/Glu, 250 mM NaCl, pH 7.5, 1 mM TCEP, 0.02% Tween, or a phosphate buffer of 140 mM KPO_4_ pH 7.4, 10 mM NaCl, 1 mM DTT, 0.02% Tween.

The FP data were gathered at room temperature using a Cytation 5 Cell Imaging Multimode reader with an excitation wavelength of 485 nm and an emission wavelength of 520 nm. The binding affinities were determined using nonlinear regression for one-site interaction using GraphPad Prism 7. The fluorescence polarization, *F*_p_, was fitted using the quadratic equation below, where the fitting parameters *F*_min_, *F*_max_, and *K*_D_ are the FP baseline, plateau, and dissociation constant, respectively. [*P*_T_] is the total protein concentration, and [*L*_T_] is the total RNA concentration (10 nM). Experiments were performed in technical triplicates.

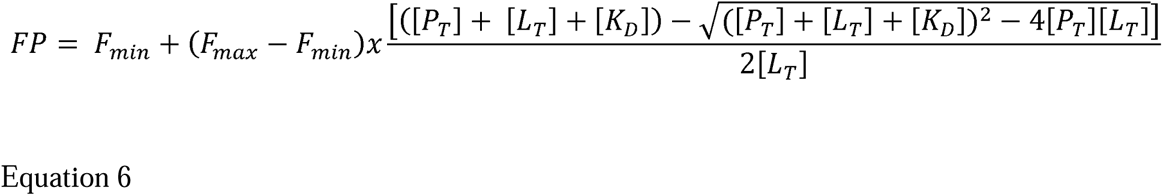

### Fluorescent imaging

SRSF1 constructs were tagged with Maleimide-Alexa488, dissolved into 800 mM Arg/Glu, pH 6.5, 0.2 mM TCEP, and stored at concentrations between 20-50 µM. Surplus Alexa488 dye was removed by a desalting column. The protein was then diluted to its final concentration into the buffer of interest and imaged on a 96-well Cellvis glass bottom plate coated with Pluronics F127. Images are brightfield/GFP channel overlays taken on a Cytation5 imager using the software Gen5 3.10. The critical concentrations were determined by imaging these dilutions at varying concentrations. The hyperphosphorylated construct was not observed to phase separate using this method at any concentration. Dialysis and buffer exchange both confirmed that the protein was soluble up to greater than 10 µM in this buffer at room temperature.

## RESULTS

### Mass spectrometry identified both the number and locations of phosphoserines on SRSF1 constructs, uncovering novel phosphorylation sites

To understand the effect of phosphorylation on SRSF1’s ensemble structures, we used various SRSF1 constructs, including its three major phosphorylation states, a construct lacking the RS tail (ΔRS) and a fourth construct termed as “pi-RS2A” (Fig. 1B). SRPK1 phosphorylates serine residues adjacent to arginine primarily on the RS1 region of the RS tail,^49,50, 51,52,26^ whereas CLK1 phosphorylates serine residues throughout the RS tail, including serines adjacent to arginine or proline.^26,53^ To restrict the placement of phosphoserines, we introduced the “pi-RS2A” construct, in which serine residues in RS2 were mutated to alanine. This left only 12 serines available for phosphorylation, all on RS1. To identify the differences between CLK1 and SRPK1 phosphorylation, the pi-RS2A construct was phosphorylated using CLK1 (Fig. 1B). We used mass spectrometry to validate the phosphorylation states of these samples prior to NMR studies (Fig. S1A, Table S1). On average, SRSF1 constructs hypo-phosphorylated by SRPK1 contained 10-13 phosphates, pi-RS2A constructs phosphorylated by CLK1 contained 9-15 phosphates, and the constructs hyperphosphorylated by CLK1 contained 18 to 23 phosphates (Fig. S1A).

The RS tail contains 16 serine residues adjacent to arginine. To test whether the 12 phosphates added by SRPK1 occur only on the RS1 region, we introduced a Lys-C cleavage site, R229K, at the boundary of RS1 and RS2 (Fig. S1B) and purified the protein in the unphosphorylated, hypophosphorylated, and hyperphosphorylated states. Previous studies have indicated that some RS2 fragment should remain unphosphorylated^26^ and that a 10:1 ratio of phosphates on RS1:RS2 should be observed.^49,54^ Consistent with the literature, after SRPK1 phosphorylation, no fragment matching the mass of unphosphorylated RS1 was found(Fig. S1D) while an unphosphorylated RS2 fragment was observed, leading to the conclusion that phosphorylation by SRPK1 preferentially occurred on the RS1 region of the RS tail. We further found that RS2 carried 0-2 phosphate groups for SRPK1-phosphorylated SRSF1 with the majority of RS2 fragments containing 1 phosphate (Fig. S1E, middle row). Because heavy phosphorylation can result in a decreased signal, we performed the same analysis on the hyperphosphorylated construct, which contains up to 20 phosphorylated serines throughout the RS tail.^26^ Hyperphosphorylation by CLK1 resulted in the addition of up to six phosphates on the RS2, corresponding to the number of serine residues in that location (Fig. S1E, bottom row). This suggests that if more heavily phosphorylated RS2 fragments were present, they would be able to be observed.

We next sought to determine if regions other than RS were phosphorylated. We observed a distribution of 18-23 phosphates for the hyperphosphorylated SRSF1 (Fig. S1C). However, only 20 serines are present in the RS tail, suggesting that regions outside of RS were also phosphorylated. It has previously been demonstrated that S119 on the RRM1/RRM2 linker can be phosphorylated by protein kinase A,^55^ although phosphorylation by CLK1 in this location has never been confirmed. To identify additional phosphorylation sites, we performed Lys-C cleavage of SRSF1 and found that, for CLK1-phosphorylated RS2A and hyperphosphorylated SRSF1, both the RRM1/RRM2 linker (Fig. S1F) and the N-terminus of the protein (Fig. S1G) were phosphorylated in some species. This finding is particularly notable because the serines on the N-terminus of the protein are adjacent to methionine and glycine residues, whereas it was previously thought that only proline or arginine residues were responsible for directing CLK1 to SRSF1’s phosphorylation sites.^53^

### Phosphorylation increases the rigidity of the RS tail

We previously obtained an NMR assignment of unphosphorylated SRSF1. For the unstructured regions that have resonance overlap, we clustered overlapping peaks to represent the regions’ chemical shift. We were able to identify spectral peaks corresponding to the arginine and serine regions of the RS tail of the phosphorylated constructs (Fig 1C, Fig. S2, Fig. S3). We analyzed the chemical shifts of SRSF1 (Fig. 1D) using TALOS, which uses a neural-network-derived algorithm to predict the order parameter (*S*^2^) and secondary structure propensity. The value of *S*^2^, which varies between 0 to 1, reflects the rigidity of bond vectors, with 0 indicating complete flexibility and 1 corresponding to complete rigidity. An order parameter of 0.85 typically indicates a fully structured region, while a secondary structure prediction of greater than 0.5 typically suggests that a region possesses a given secondary structure. Thus, anything below 0.5 is likely unstructured. To complement these data, we obtained ^15^N transverse relaxation time constant (*T_2_*). *T_2_* values are sensitive to molecular tumbling and local structural dynamics. Unstructured regions such as N- and C-termini have higher flexibility and therefore a larger *T_2_*, compared with structured regions. We found robust agreement between the known secondary structures of the RRM domains and the TALOS predictions (Fig. 1D). *T_2_* relaxation and TALOS-derived parameters indicated that the RRM1/RRM2 linker and RS were highly flexible. However, we observed some residues in this region having relatively high *S*^2^ and secondary structure propensities, which indicates a possibility that transient secondary structure may occur. Upon hyper-phosphorylation, the *S*^2^ of RS increases across the entire region, indicating an increase in the rigidity (Fig. 1D, middle panel). Consistent with the order parameter analysis, hyper-phosphorylation decreases the *T*_2_ values of SRSF1 (Fig. 1D, top panel). This is consistent with previous studies on the effect of phosphorylation on isolated RS domains.^56^ Interestingly, pi-RS2A, which has phosphorylated RS1 and unphosphorylated RS2, demonstrates segmental *T*_2_ values, with RS1 resembling hyper-phosphorylated SRSF1 and RS2 resembling the unphosphorylated protein (Fig. 1D, top panel).

### Phosphorylation decreases interactions between the RS tail and RRM1 helices in favor of interactions with the RRM1 RNA-binding site

To determine how phosphorylation of the RS tail altered interactions with the other portions of SRSF1, we analyzed the chemical shift perturbations (CSPs) between different full-length SRSF1 constructs and the ΔRS construct (Fig. S4). A CSP above the standard deviation of the spectrum is indicative of changes in the immediate chemical environment. This can be due to direct formation of bonds, breaking of bonds, or conformational changes. Regardless of the phosphorylation state, introduction of an RS tail caused significant CSPs across RRM2 (Fig. S4A, S4B). As phosphorylation increased, slightly more interactions were observed in positively charged regions adjacent to RNA-binding residues—namely, the loop adjacent to the β3 sheet of RRM1 and the first α-helix of RRM2 (Fig. S4A). Decreasing the salt concentration enabled more CSPs to be observed. Namely, in 200 mM Arg/Glu (Fig. S4B), interactions of the unphosphorylated RS tail with the helices of the RRM1 domain became more pronounced when arginine was further reduced, and phosphate ions were present (Fig. S4C). To directly characterize these interactions, we used paramagnetic relaxation enhancement (PRE) NMR. PRE complements CSPs by providing transient long-range distance restraints. A higher PRE indicates that the labeling site comes closer to the residue of interest, following a distance relationship of r^− 6^.^57^ Bleaching of the signal indicates the PRE was too large for accurate determination, suggesting that the probe comes within a distance of 12 Å (see methods section for how distances were averaged).^57^ We placed the paramagnetic probe at N220C at the center of the tail (Fig. 2), or T248C at the C-terminal end (Fig. S4D). Our PRE data suggested that unphosphorylated RS transiently interacted with residues around the α1 helix of the RRM1 domain, while phosphorylated RS had more interactions with RNA-binding residues (Fig. 2B, 2C). The hyperphosphorylated construct of SRSF1 is soluble in a phosphate buffer while the other constructs are not. This provided us with the opportunity to perform experiments in the presence of a relatively low salt phosphate buffer of 140 mM KPO_4_ and 10 mM NaCl. The iPRE pattern matches a similar trend to the hyperphosphorylated PRE data (Fig. 2B). However, we observed more interactions between RS and the RNA-binding sites, such as β1 of RRM1. suggesting that the buffer composition modulates the strength, but not the location, of the interactions.

**Figure 2.**
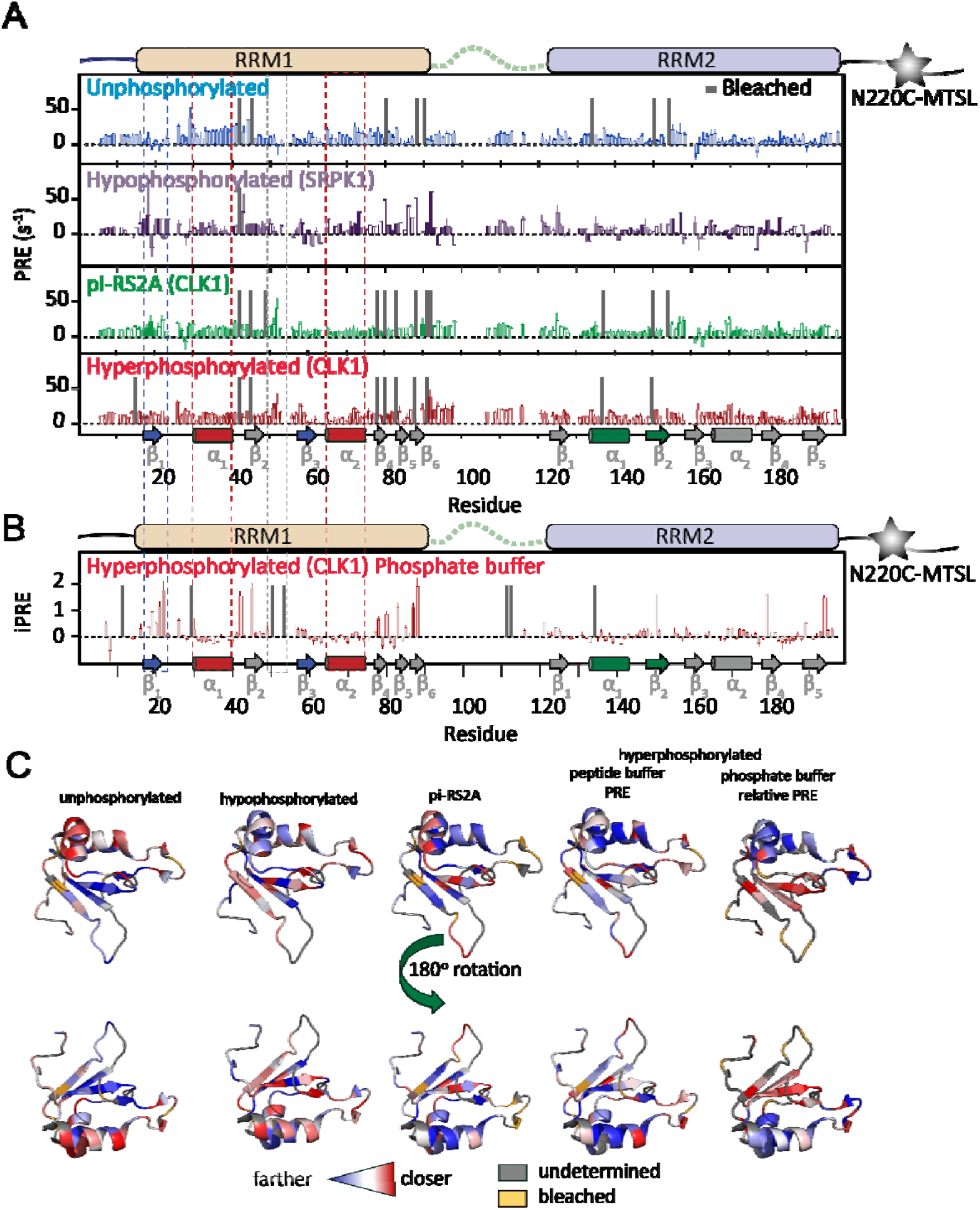
Phosphorylation of the RS tail decreases its interactions with the RRM1 helices and increases its interactions with residues on or adjacent to the RNA-binding site. (A) PRE of SRSF1 in its four major phosphorylation states with a tag placed on residue N220C in a buffer of 100 mM ER4, 400 mM Arg/Glu, pH 6.3. Secondary structures are presented at the bottom of the plot. RRM1 is shown in blue while the helices responsible for protein-protein interactions are shown in red. RNA-binding residues of RRM2 are shown in green. Phosphorylation results in fewer interactions with the RRM1 helices (red dashes) and more interactions with RNA-binding residues (blue dashes) and adjacent loops (grey dashes). (B) iPRE of hyperphosphorylated SRSF1 in 140 mM KPO_4_ pH 7.4, 10 mM NaCl, with a tag placed on residue N220C adjacent to PyMOL projection. The preference for RNA-binding residues on RRM1 and lack of preference for the helices is exacerbated under these conditions. iPRE values were calculated as the logarithm of the intensity ratio of dia- to para-magnetic states. (C) PyMOL projections of PRE values onto the solution structure of RRM1 (PDB ID: 6HPJ).

### Phosphorylation changes driving forces of SRSF1 LLPS

Our previous study has shown that the RS tail of SRSF1 plays a critical role in LLPS of SRSF1 in vitro.^25^ In this study, we found that the LLPS mechanisms by which SRSF1 phase separates depends on its phosphorylation state. In phosphate buffer, the critical concentration for SRSF1 LLPS increases from approximately 0.1 µM for unphosphorylated SRSF1 to approximately 4 µM for pi-RS2A and to larger than 10 µM for hyper-phosphorylated SRSF1 (Fig. 3A).

**Figure 3.**
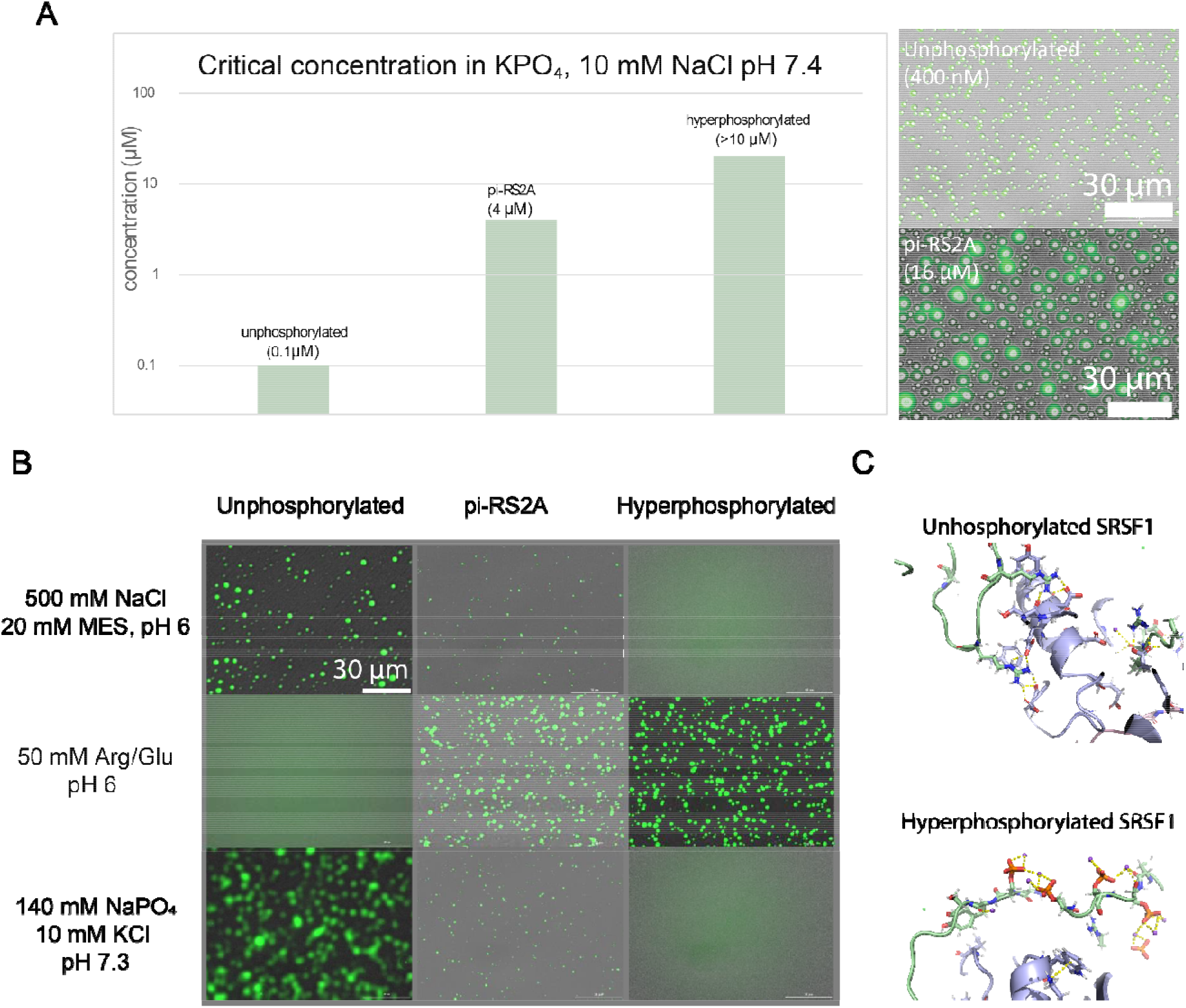
SRSF1 constructs phase separate under distinct conditions. (A) Critical point determination of unphosphorylated and pi-RS2A in a potassium phosphate buffer of 140 mM KPO_4_ pH 7.4, 10 mM NaCl, at room temperature, adjacent to sample images of SRSF1 diluted into phosphate buffer at indicated concentrations. (B) Phase separation of SRSF1’s various phosphorylation states in various buffers at 300 nM. Proteins are tagged with Alexa488 at residue T220C. Images are GFP/brightfield channel overlays. (C) MD simulations of SRSF1 in the presence of 150 mM NaCl in distinct phosphorylation states. The hyperphosphorylated RS domain is predicted to form extensive interactions with sodium, which may prevent intermolecular contacts. The unphosphorylated tail forms more stable interactions with aromatic regions of the RRM domains, with which ionic salts cannot compete well.

Our and others’ studies have indicated that insight into the interactions driving LLPS can be obtained by analyzing the buffers best suited for solubilizing the protein.^25, 58^ To better understand the effect of phosphorylation on LLPS, we diluted fluorescently labeled SRSF1 into various buffers and visualized their LLPS through fluorescent imaging. In a potassium phosphate buffer or a buffer containing 500 mM NaCl, the tendency for SRSF1 LLPS followed the trend of: unphosphorylated > pi-RS2A > hyper-phosphorylation. However, the trend in LLPS tendency was reversed in 50 mM Arg/Glu (Fig. 3B). We reasoned that the reversal is due to a change in the driving force for liquid-liquid phase separation upon phosphorylation. In the unphosphorylated state, Arg residues in the RS tail play a dominant role in phase separation. Therefore, Arg amino acids in the buffer best compete for Arg-mediated intermolecular interactions. In contrast, ionic salts such as Na^+^, or phosphate ions cause salting out, consistent with Arg’s ability to form cation-pi stacking and hydrophobic interactions in addition to electrostatic interactions.^58^ This salting out is similar to previous observations on proteins that phase separate through Arg-driven hydrophobic interactions.^59^ In the phosphorylated state, phosphorylated Ser residues also play a role in LLPS. The interactions mediated by phosphoserine are electrostatic. Therefore, ions like Na^+^, K^+^ or phosphate can efficiently inhibit the intermolecular interaction responsible for LLPS. Consistent with our proposed rationale, our MD simulation observed hyperphosphorylated RS has extensive interactions with Na^+^ while unphosphorylated RS has more interactions with aromatic or acidic residues (Fig. 3C).

### Phosphorylation of the RS tail changes the interactions between the RRM domains and the RRM1/RRM2 linker

SRSF1 has a 30-amino acid unstructured linker between its two RRM domains. The RRM1/RRM2 linker contains Arg-rich sequences on both N- and C-terminal ends (Fig. 4A). Although SRPK1 and CLK1 are known to phosphorylate RS of SRSF1, we see evidence that the phosphorylation of the RS tail also affects the behavior of the RRM1/RRM2 linker. Chemical shift and *T_2_* relaxation data suggested that the linker becomes more rigid upon phosphorylation (Fig. 1D). To identify the intramolecular interactions mediated by the linker, we compared the chemical shifts of the ΔRS construct with the individual RRM domains. By comparing the ΔRS construct and RRM1 (Fig. 4A, left panel), we found that most perturbations were clustered on the RRM1 α2 helix. By comparing the ΔRS construct and RRM2 (Fig. 4A, right panel), CSPs were found on the RRM2 β2 and β3 sheets as well as the N-terminal portion of the α2 helix. Interestingly, when the C-terminal portion of the linker was attached to RRM2, its CSPs against the ΔRS construct were abolished (Fig. 4B, right), suggesting that this portion of the linker is responsible or able to compensate for the linker-involved intramolecular interactions. On the other hand, when the N-terminal portion of the linker was attached to RRM1, its CSP pattern (Fig. 4B, left) is similar to that of RRM1 alone (Fig. 4A, left), suggesting that the N-terminal portion of the linker is not responsible for the linker-mediated interactions with RRM1. These results together suggested that the C-terminal portion of the RRM1/RRM2 linker was important to interactions between the two RRM domains and the linker. To gain insight into how phosphorylation of the RS tail affects the interaction between the linker and other domain in SRSF1, we placed a paramagnetic tag at the end of the linker (E120C) and collected PRE data in different phosphorylation states (Fig. 4C). Due to the long perturbation range of the paramagnetic group, several RRM1 and RRM2 regions were bleached. Interestingly, we found that upon phosphorylation, PRE on the RRM1 α2-helix was significantly reduced (Fig. 4C). We noted that residues showing high PRE or bleaching (Fig. 4D) resembled those with high CSPs (Fig. 4A, 4B), especially the α2 helix and neighboring loops of RRM1. Therefore, our CSP and PRE data together suggests that the C-terminal portion of the RRM1/RRM2 linker interacts with RRM1 and RRM2, and phosphorylation of RS weakens these linker-mediated intramolecular interactions.

**Figure 4:**
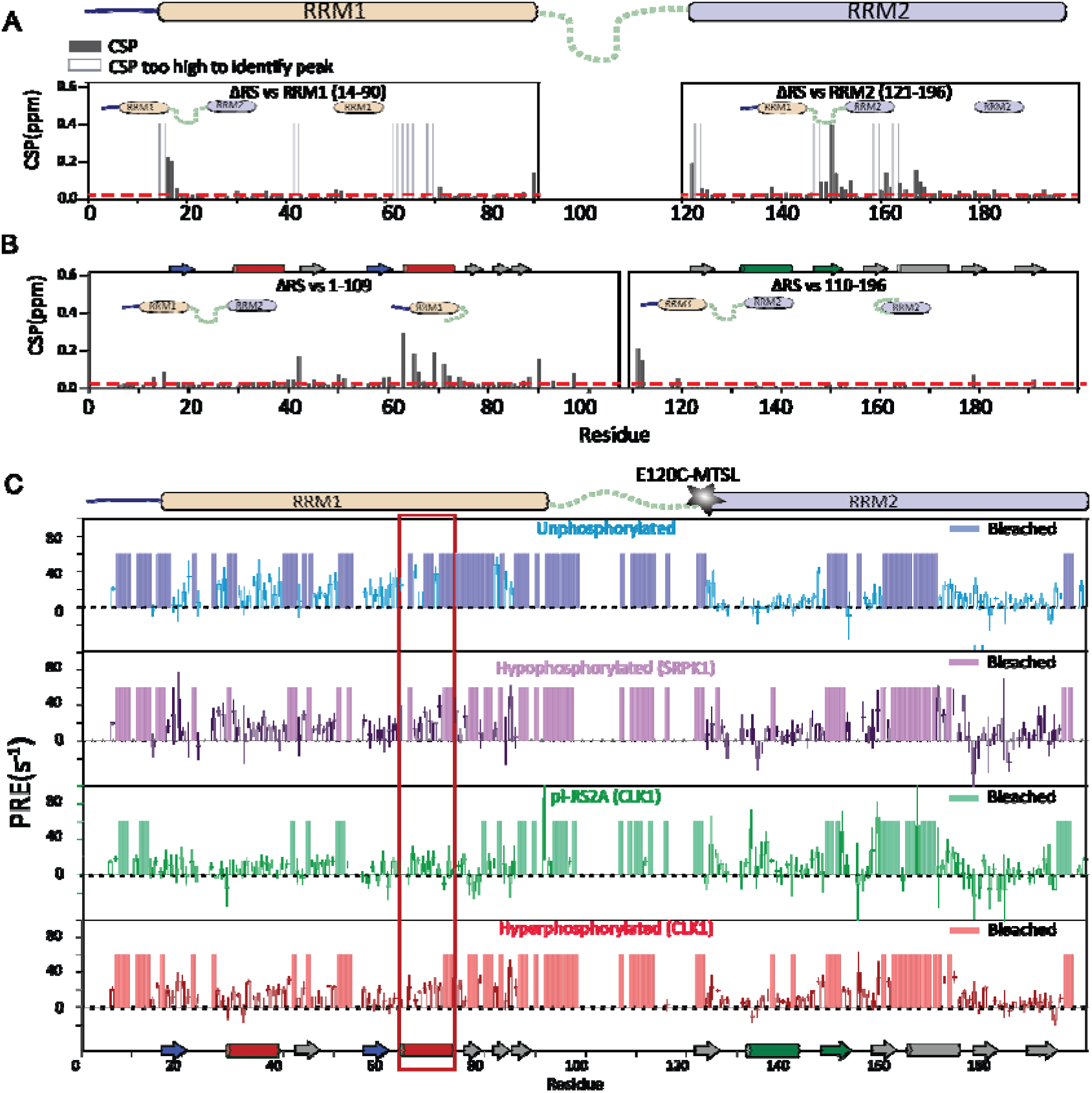
The α2 helix of RRM1 interacts with the interface between the RRM1/RRM2 linker and RRM2, and phosphorylation decreases these interactions. (A) CSPs between individual RRM domains and the ΔRS construct. (B) CSPs between the ΔRS construct and RRM domains containing segments of the RRM1/RRM2 linker. (C) PRE of SRSF1 in four phosphorylation states with a tag placed on residue E120C. The α2 helix location is highlighted by a red box.

### Phosphorylation inhibits the RNA binding of SRSF1

Our PRE and CSP data revealed that the phosphorylated RS tail interacted with the RNA-binding site of RRM1. These results hint the inhibitory effect of the phosphorylated RS tail on the RNA binding of SRSF1. The RRM domains of SRSF1 have distinct RNA-binding preferences. RRM1 recognizes cytosine residues preferentially, whereas the RRM2 domain of this protein recognizes purine residues, such as motifs “GGA” and “GGG.” Neither domain recognizes uracil residues with specificity. Because an RNA ligand lacking both RRM1 and RRM2 cognate motifs binds poorly to SRSF1 regardless of phosphorylation state, we used semi-target RNA lacking either the RRM1 or RRM2 cognate recognition motif (Fig. 5A). Compared to a ΔRS ligand, unphosphorylated SRSF1 binds more tightly to both targets (K_D_ = 38 ± 2 nM vs 104 ± 3 nM) (Fig. 5B) and semi-target (K_D_ = 400 ± 20 nM vs 2400 ± 100 nM) (Fig. 5C) ligands. Phosphorylation weakens the RNA binding of SRSF1 (Fig. 5B, 5C). In the phosphate buffer, the tendency of the phosphorylated RS to decrease binding affinity becomes more pronounced, with hyperphosphorylation resulting in binding weaker than the ΔRS construct. This was the case for both target and nontarget RNA, indicating that the phosphorylated RS tail inhibited both RRM1 and RRM2. This is consistent with the fact that the phosphorylated tail interacts with both binding sites (Fig. 2).

**Figure 5.**
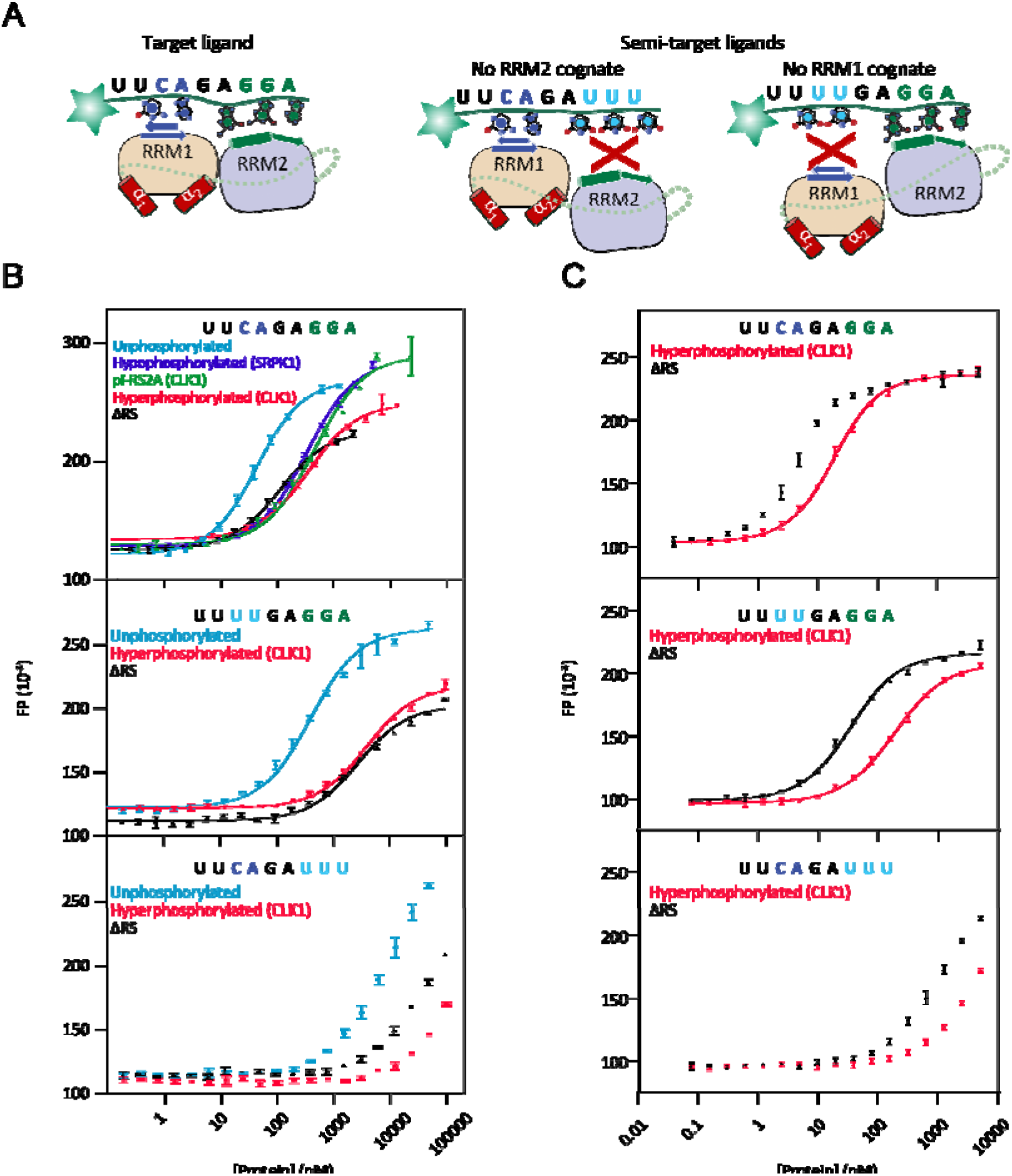
Binding to target and non-target RNA is weakened by phosphorylation, and this trend becomes more pronounced in phosphate buffer. (A) RNA ligands used in these experiments. Fluorescein tag was placed at the 5’ end of the RNA (indicated by a star). (B) Binding of various constructs of SRSF1 to its target RNA ligand. An arginine buffer was used consisting of 250 mM Arg/Glu, 250 mM NaCl, 10 mM HEPES pH 7.5, 0.1 mM TCEP, 0.02% NaN_3_ (C) Binding in this arginine buffer of SRSF1 constructs to target and semi-target RNA. (D) Binding to target and semi-target RNA in a phosphate buffer of 140 mM KPO_4_ pH 7.4, 10 mM NaCl, 1 mM DTT, 0.02% Tween.

### Phosphorylation of SRSF1 increases surface exposure of the RRM1 helices and the interactions between unstructured regions

SRSF1 exists as a dynamic ensemble of fast exchanging conformers due to the flexibility of the RS tail and the RRM1/RRM2 linker. However, our CSP and PRE data also indicate that intramolecular interactions within SRSF1 are not random. To gain deeper insight into the intramolecular interactions and their regulation by phosphorylation, we simulated the ensemble structure of SRSF1 using the experimental CSP and PRE data. With the distance restraints derived from these experimental data, we first built the model using Xplor-NIH, followed by AMBER20 molecular dynamics to refine the chemical bonds/angles and local interactions. As illustrated in Fig. 6A, an increased surface accessibility of the RRM1 helices upon phosphorylation was observed. We next analyzed the types of intramolecular interactions within the SRSF1 ensemble. We found various stacking interactions such as pi-pi stacking, Arg-Arg stacking, and cation-pi stacking interactions mediated by arginine, as well as H-bonds and salt bridges. In the unphosphorylated state, extensive cation-pi stacking occurred between the unphosphorylated RS tail and the RRM domains as well as within the unstructured regions themselves (Fig. 6B). Meanwhile, H-bonds/salt bridges occurred between the RS tail and RRM1 helices (Fig. 6B). In the hypophosphorylated state, stacking and H-bonds/salt bridges to the RRM1 helices still existed, particularly with the C-terminal portion of the tail, which was still unphosphorylated (Fig. 6C, and 6D). However, fewer stacking interactions were observed overall, and there was an increase in H-bonds/salt bridges both within and between the unstructured regions (Fig. 6D). A similar trend was observed for the pi-RS2A construct, although more cation-pi interactions were observed with RRM2, and more extensive Arg-Arg and H-bonds/salt bridges within the tail were formed for the pi-RS2A construct (Fig. 6C). Finally, in the phosphorylated state, interactions with the RRM1 domain primarily involved H-bonding with the β1, β3 sheets, and neighboring loops (Fig. 6E, 6F). Moreover, extensive H-bonds/salt bridges were found to occur within the RS tail (Fig. 6E, 6F).

**Figure 6.**
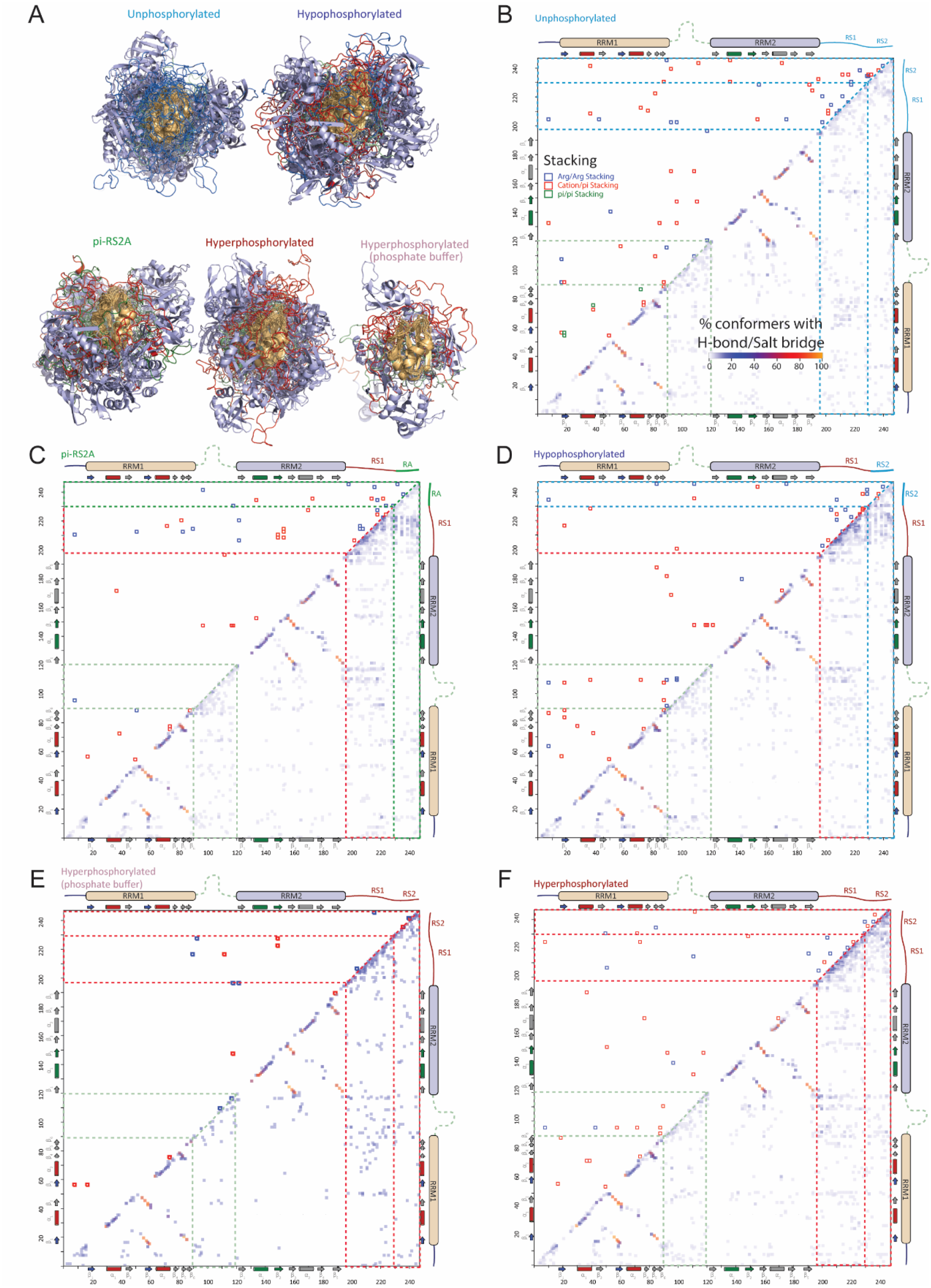
Phosphorylation increases surface exposure of the RRM1 helices and interactions within RS and between RS and the RRM1/RRM2 linker. (A) Conformer overlays of the SRSF1 ensemble in the five phosphorylation states. Conformers were aligned using RRM1 (golden) as the anchor domain. RRM2 and RS were shown in light blue cartoon and ribbon, respectively. The un-phosphorylated and phosphorylated segments of RS were shown in green and red, respectively. The color scheme is the same as the domain architecture in the following panels. As the protein becomes phosphorylated, the RRM1 helices become more surface-exposed. (B-F) Heatmap analysis of intramolecular interactions within SRSF1 in different phosphorylated states. Stacking interactions, such as Arg/Arg, cation/pi, pi/pi, are shown above the diagonal by open squares. H-bonding/salt bridge interactions are shown below the diagonal by filled squares with the color scale showing its occurrence in percentage.

### Autoinhibition is a possible mechanism by which phosphorylation alters SRSF1’s binding preferences

In addition to RNA-binding sites, the RS-mediated sites on SRSF1 also include the two RRM1 α-helices, which are responsible for interacting with U1-70K and PP1.^22–24^ Binding to U1-70K follows a trend opposite to that of RNA: it is tightest when SRSF1 is hyperphosphorylated and loosest when SRSF1 is unphosphorylated.^24^ PP1 follows a trend similar to U1-70K, but SRSF1’s RRM1 domain is crucial for mediating the effect of phosphorylation state on PP1 binding. SRSF1’s RRM1 domain tightens PP1’s initial binding to hyperphosphorylated SRSF1, but the same RRM1 domain also weakens PP1’s binding to dephosphorylated constructs, ensuring dephosphorylation stops before SRSF1 is completely unphosphorylated^23, 60, 61^. Moreover, phosphorylation of RS alters the intramolecular interactions.

Therefore, analyzing the relationship between intramolecular interactions within SRSF1 and its RNA/protein binding sheds light on these important events during the spliceosome assembly. To this end, we compared the SRSF1 ensemble and its complex with RNA, U1-70K, or PP1. In the phosphorylated state, RS interacts with the RRM1 arginine residues involved in RNA binding (Fig. 7A, 7B). This explains the inhibitory effect of RS on RNA binding in the phosphorylated state.

**Figure 7.**
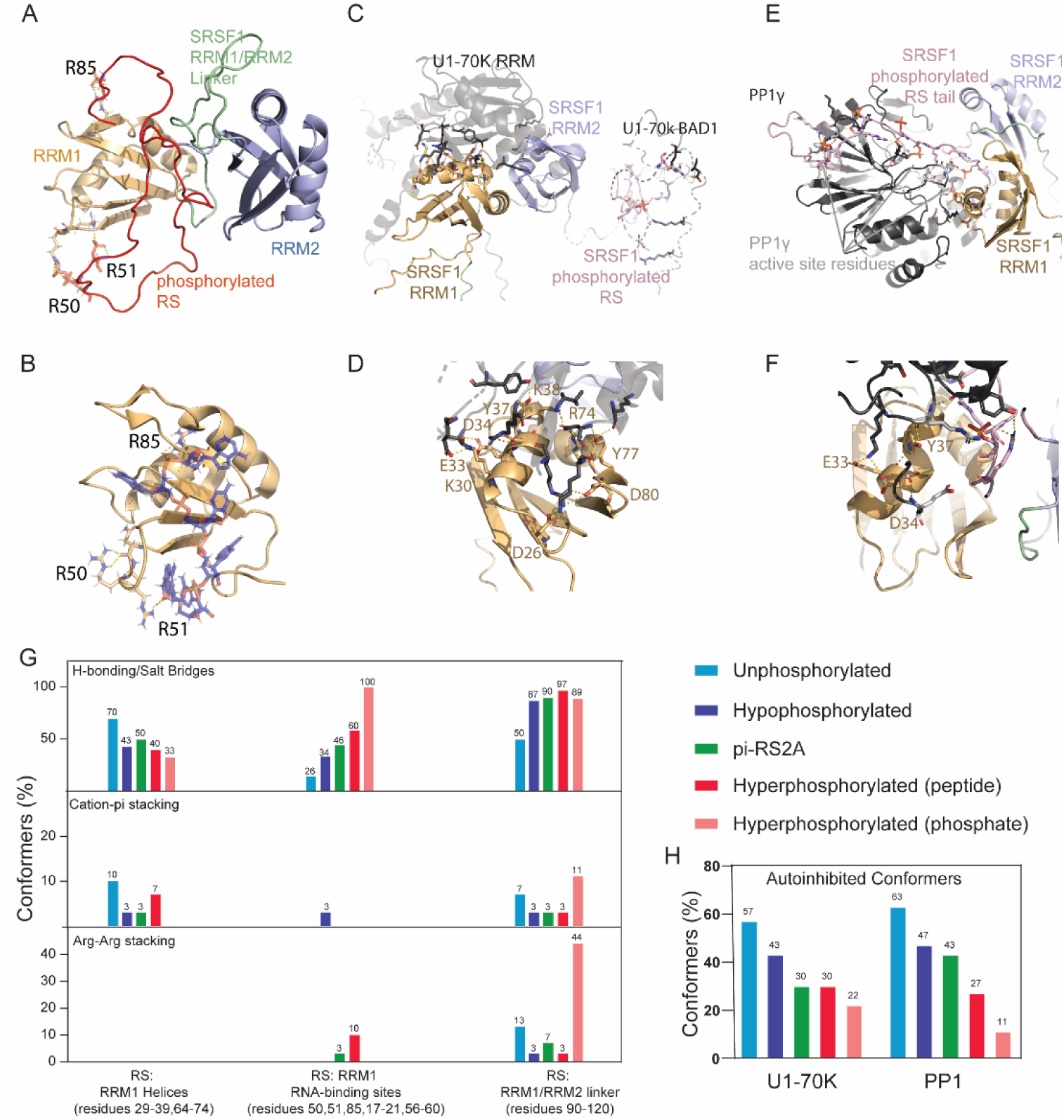
Phosphorylation regulates autoinhibition of SRSF1 by altering binding of SRSF1 to protein and RNA binding partners. (A) A representative conformer from the phosphorylated SRSF1 ensemble demonstrating the interaction between phosphorylated RS and the RNA-binding sites on RRM1 and RRM2. (B) The RNA-bound structure (PDB ID: 6HPJ) highlights the arginine residues involved in RNA (sticks) binding. Note that these arginine residues are inhibited by phosphorylated RS (red ribbon in panel A). (C) AlphaFold3-predicted SRSF1/U1-70K complex. The interactions revealed by the predicted complex are consistent with our previous experimental discoveries: SRSF1 RRM1’s helices interact with U1-70K RRM, phosphorylated SRSF1 RS interacts with U1-70K BAD domains. (D) Panel C expanded. (E) AlphaFold3-predicted SRSF1/PP1 interactions. (F) Panel E expanded. (G) Occurrence of intramolecular interactions between the SRSF1 RS tail (Residues 197-248) and RRM1 helices, RRM1 RNA-binding sites, and the RRM1/RRM2 linker, in different phosphorylation states. Y axis indicates percentage of conformers in an ensemble structure with a given interaction. (H) Percentage of autoinhibited conformers coming within a 4 Å surface of the binding partner.

The high-resolution structures have not yet been obtained for SRSF1 in complex with either U1-70K or PP1. However, the regions of SRSF1 involved in interactions with these proteins have been determined^23, 24, 62^. We used AlphaFold3 to predict the structure for SRSF1 bound to U1-70K or PP1. The predicted structure is consistent with our previous findings that the phosphorylated SRSF1 RS interacts with U1-70K BAD1 domain while SRSF1 RRM1 interacts with a loop region of U1-70K RRM. (Fig. 7C, 7D).^24^ Using AlphaFold3, we also predicted the SRSF1/PP1 complex structure, which reveals a PP1-binding pocket between the helices of SRSF1’s RRM1 domain (Fig. 7E, 7F) and SRSF1 RS interacting with the PP1 active site for dephosphorylation.^23, 62–64^ These features of the predicted SRSF1/PP1 complex structure also agree with previous reported experimental studies.^23^

We further analyzed how phosphorylation regulates the intramolecular interactions mediated by the RS tail (Fig. 7G). We found that progressive phosphorylation resulted in increased interactions with the RNA-binding sites but decreased interactions with the two RRM1 helices. These trends are in line with the observations that phosphorylation inhibits RNA binding but promotes interacting with U1-70K or PP1. In addition, we also noticed that some conformers in the SRSF1 ensemble formed intramolecular interactions that would prevent binding to U1-70K or PP1 complex. We defined an autoinhibited conformer as a conformer in which the binding site of a given partner was occluded. We identified autoinhibited conformers by aligning them with the RRM1 domain in the AlphaFold structure and determining whether the remainder of the protein overlapped with the surface of either PP1 or U1-70K. Interestingly, the occurrence of conformers autoinhibited against either U1-70K or PP1 decreased dramatically with the progression of RS tail phosphorylation (Fig. 7H). This trend suggests that in the phosphorylated state, the entropy penalty for U1-70K or PP1 binding is reduced, facilitating the binding process.^65, 66^

## DISCUSSION

### Summary of Key Findings

In this study, we systematically characterized the intramolecular interactions within SRSF1 in its unphosphorylated, hypophosphorylated, and hyperphosphorylated states. Our key findings include: (1) in the unphosphorylated state, the RS tail primarily interacts with electronegative patches on the α1 and α2 helices of RRM1; (2) as phosphorylation progresses, the RS domain shifts its interaction to the RNA-binding sites on RRM1 and RRM2; (3) in the phosphorylated state, arginine residues in the RS tail form salt bridges with neighboring phosphorylated serine residues; (4) CLK1 phosphorylates both the N-terminus and the RRM1/RRM2 linker of SRSF1, while SRPK1 adds 1-2 phosphate groups to the C-terminal half of the RS tail (RS2).

### Coordination of RNA binding and protein interactions through SRSF1 phosphorylation in early spliceosome assembly

Association of SRSF1 with mRNA starts at the transcription step via interactions with RNA polymerase II.^2, 3^ Previous studies have shown that the RS tail directly contacts mRNA.^67, 68^ Our findings, along with those of others, demonstrate that unphosphorylated RS significantly enhances the RNA-binding affinity of SRSF1.^69^ This exceptionally high affinity could potentially lead to cryptic RNA binding. However, erroneous splicing is unlikely, as the α1 and α2 sites of RRM1, which interact with U1-70K, are blocked by the unphosphorylated RS domain. As phosphorylation increases, the inhibitory effect of RS ensures that only authentic exonic splicing sites persist under this competitive pressure. Meanwhile, phosphorylated RS gains the ability to interact with the basic-acidic dipeptide domain 1 (BAD1) of U1-70K, as shown in our previous study.^24^ Additionally, the U1-70K binding sites on SRSF1’s RRM1 becomes accessible, enhancing U1-70K recruitment to authentic splicing sites.^24^ Upon formation of the complex with U1-70K, the RS tail of SRSF1 engages in interactions with U1-70K’s BAD1 domain, relieving the inhibition on RNA binding. This is consistent with previous findings showing that the RNA: SRSF1 complex is more stable.^22^

It has been reported that activation of the spliceosome requires partial dephosphorylation of SRSF1, which is mediated by PP1.^29^ Before the spliceosome can become activated, the U1 complex must also dissociate from the spliceosome. Interestingly, U1-70K and PP1 share a common binding site between the α1 and α2 helices of RRM1, which, as we demonstrate, is involved in intramolecular interactions with the unphosphorylated RS tail.^24^ Therefore, it is plausible that PP1 competes with U1-70K for this site on RRM1, leading to RS dephosphorylation, which weakens the interaction between SRSF1 and U1-70K and promotes the dissociation of U1 snRNP from the spliceosome.

### Hypophosphorylated and dephosphorylated SRSF1 may behave differently

The differences we see between the PRE patterns of pi-RS2A phosphorylated by CLK1 and SRSF1 that has been hypophosphorylated by SRPK1 suggest that number of phosphates is not the only factor in the differences between the roles of SRSF1 in its hyper vs hypophosphorylated states. Differences we observe in the location, rather than just the number, of phosphate residues added might be a driving factor in this (Fig. S1). Evidence suggests that while PP1 dephosphorylates the C-terminus with preference over the N-terminus, it likely does not reproduce the original hypophosphorylated state exactly.^70^ As it remains to be explored whether phosphatases can dephosphorylate the linker and N-terminus, SRSF1 in its post-splicing, dephosphorylated state may be capable of serving subtly distinct roles from the hypophosphorylated protein.

### Regulation of SRSF1 phase separation by phosphorylation

LLPS is mediated by multivalent weak interactions, including electrostatic, polar, H-bonding, pi-pi stacking, cation-pi stacking, and hydrophobic interactions. Of all amino acid residues, arginine is unique due to its ability to form all aforementioned weak interactions. It is noteworthy that Arg-Arg stacking driven by hydrophobicity has a considerable strength. A recent study has found that Arginine-rich peptides, deprived of aromatic residues or acidic residues, are able to form LLPS.^59^ Obviously, the hydrophobic interaction among arginine residues outperforms the electrostatic repulsion. This versatility explains why Arg-rich regions are responsible for phase separation of many proteins, such as hnRNPA1 and FUS.^71–74^

Our previous work has demonstrated that the RS domain playing a major role in SRSF1 LLPS.^25^ We found that short peptides mimicking the RS tail can efficiently solubilize SRSF1 (up to 26-fold more efficiently than Arg/Glu in the same concentration), but peptides that do not accurately mimic the RS tail are less effective at solubilizing SRSF1 (up to 2-fold more efficient than Arg/Glu in the same concentration). In this study, we show that unphosphorylated and phosphorylated demonstrate distinct phase separation behavior. Arginine buffers are more competent to dissolve unphosphorylated SRSF1 while phosphate or salt buffers are more competent to dissolve phosphorylated SRSF1. These results together suggest that arginine residues play a dominant role in LLPS of unphosphorylated SRSF1, while phosphorylated SRSF1 LLPS depends more on phosphoserine residues. Our experimental-guided MD simulations revealed that phosphorylation shifts the RS-mediated interactions from interdomain to intradomain (Fig. 6). We find that arginine residues are engaged in hydrogen bonds or salt bridges with nearby phosphoserine residues. These intradomain interactions reduce the availability of arginine residues for intermolecular interactions. Therefore, phosphorylation of RS reduces LLPS.

Beyond SR proteins, SR-related proteins, comprising 40-50 members, such as U1-70K, SON, and SRRM2, also contain arginine/serine dipeptide repeats.^75–77^ Many of these proteins have been found to subject to phosphorylation in the cell, suggesting that the regulative mechanism by phosphorylation could be of generality.^75–77^ Therefore, the effects of phosphorylation on intramolecular interactions revealed in this study likely extend to other SR and SR-related proteins.

**Table 1:** Dissociation constants (K_D_) of SRSF1 binding to target and semi-target RNA in Arginine/Glutamic Acid (corresponds to Fig. 5B)

| RNA probe<br>(10 nM) | SRSF1 titrant | $K_D$ (nM) | RRM1<br>Selectivity for<br>CA/UU |
| --- | --- | --- | --- |
| UUCAGAGGA | Unphosphorylated | $38 \pm 2$ | -- |
| UUCAGAGGA | Hypophosphorylated | $320 \pm 10$ | -- |
| UUCAGAGGA | pi-RS2A | $500 \pm 40$ | -- |
| UUCAGAGGA | Hyperphosphorylated | $340 \pm 20$ | -- |
| UUCAGAGGA | ΔRS | $104 \pm 3$ | -- |
| UUUUGAGGA | Unphosphorylated | $400 \pm 20$ | $10.5 \pm 0.7$ |
| UUUUGAGGA | Hyperphosphorylated | $6000 \pm 300$ | $18 \pm 1$ |
| UUUUGAGGA | ΔRS | $2400 \pm 100$ | $23 \pm 1$ |
| UUCAGAUUU | Unphosphorylated | N.D. | -- |
| UUCAGAUUU | Hyperphosphorylated | N.D. | -- |
| UUCAGAUUU | ΔRS | N.D. | -- |

\*N.D. denotes binding too weak to fit accurately using these conditions

**Table 2:** Dissociation constants (K_D_) of SRSF1 binding to target and semi-target RNA in phosphate buffer (corresponds to Fig. 5C)

| RNA probe (10 nM) | SRSF1 titrant | $K_D$ (nM) | RRM1<br>Selectivity for<br>CA/UU |
| --- | --- | --- | --- |
| UUCAGAGGA | Hyperphosphorylated | $13.7 \pm 0.5$ | -- |
| UUCAGAGGA | ΔRS | Below detection limit<br>($<10$) | -- |
| UUUUGAGGA | Hyperphosphorylated | $181 \pm 6$ | $13.2 \pm 0.7$ |
| UUUUGAGGA | ΔRS | $30 \pm 2$ | -- |
| UUCAGAUUU | Hyperphosphorylated | N.D. | -- |
| UUCAGAUUU | ΔRS | N.D. | -- |

\*N.D. denotes binding too weak to fit accurately using these conditions

## Acknowledgements

We want to thank the manager of UAB Central Alabama High-Field NMR Facility Dr. Ron Shin, the director of the NMR facility Dr. William Placzek, the director of UAB Structural Biology Core Facility Dr. Champion Deivanayagam, Dr. Charles D. Schwieters at the NIH for technical support. This work was supported by National Science Foundation (MCB 2024964 to Jun Zhang).

