## Supplemental for "Controlled by disorder: phosphorylation modulates SRSF1 domain availability for spliceosome maturation"

| Construct | Hypophosphorylated (SRPK1) | pi-RS2A (CLK1) | Hyperphosphorylated (CLK1) |
| --- | --- | --- | --- |
| ^15^N SRSF1 C16/148S **N220C** | ${12}_{-2}^{+1}$ | ${12}_{-4}^{+3}$ | -- |
| ^15^N SRSF1 C16/148S **E120C** | ${12}_{-2}^{+1}$ | ${12}_{-4}^{+3}$ | ${20}_{-7}^{+3}$ |
| ^15^N SRSF1 C16/148S **T248C** | ${12}_{-2}^{+1}$ | ${12}_{-1}^{+2}$ | ${19}_{-3}^{+3}$ |
| WT | ${11}_{-1}^{+1}$ | ${11}_{-3}^{+2}$ | -- |

Table S1: Phosphates added by construct

Where major species = M, lowest value observed= l, highest value observed = u: $M_{-l}^{+u}$

Table S2: Fitting values for Xplor-NIH structure calculations.

| Construct | Labeling site | Structured Region | Q (PRE) | R^2^ (PRE) | Artificial NOE violations |
| --- | --- | --- | --- | --- | --- |
| Unphosphorylated  (30 conformers) | 120C | RRM1 | 0.299 | 0.817 | 1 |
|  | 220C | RRM1 | 0.218 | 0.985 |  |
|  | 220C | RRM2 | 0.326 | 0.908 |  |
|  | 248C | RRM1 | 0.283 | 0.996 |  |
|  | 248C | RRM2 | 0.228 | 0.983 |  |
| Hypophosphorylated  (30 conformers) | 120C | RRM1 | 0.345 | 0.962 | 0.2 |
|  | 220C | RRM1 | 0.372 | 0.961 |  |
|  | 220C | RRM2 | 0.287 | 0.959 |  |
|  | 248C | RRM1 | 0.318 | 0.916 |  |
|  | 248C | RRM2 | 0.249 | 0.974 |  |
| pi-RS2A  (30 conformers) | 120C | RRM1 | 0.312 | 0.999 | 0.2 |
|  | 220C | RRM1 | 0.226 | 0.945 |  |
|  | 220C | RRM2 | 0.148 | 0.975 |  |
|  | 248C | RRM1 | 0.400 | 1.000 |  |
|  | 248C | RRM2 | 0.203 | 0.965 |  |
| Hyperphosphorylated  (30 conformers) | 120C | RRM1 | 0.236 | 0.998 | 0.1 |
|  | 220C | RRM1 | 0.290 | 0.997 |  |
|  | 220C | RRM2 | 0.265 | 0.993 |  |
|  | 248C | RRM1 | 0.282 | 0.965 |  |
|  | 248C | RRM2 | 0.401 | 0.920 |  |
| Hyperphosphorylated (iPRE in phosphate buffer)  (9 conformers) | 220C | RRM1 | -- | 0.948 | 0.2 |
|  | 220C | RRM2 | -- | 0.948 |  |

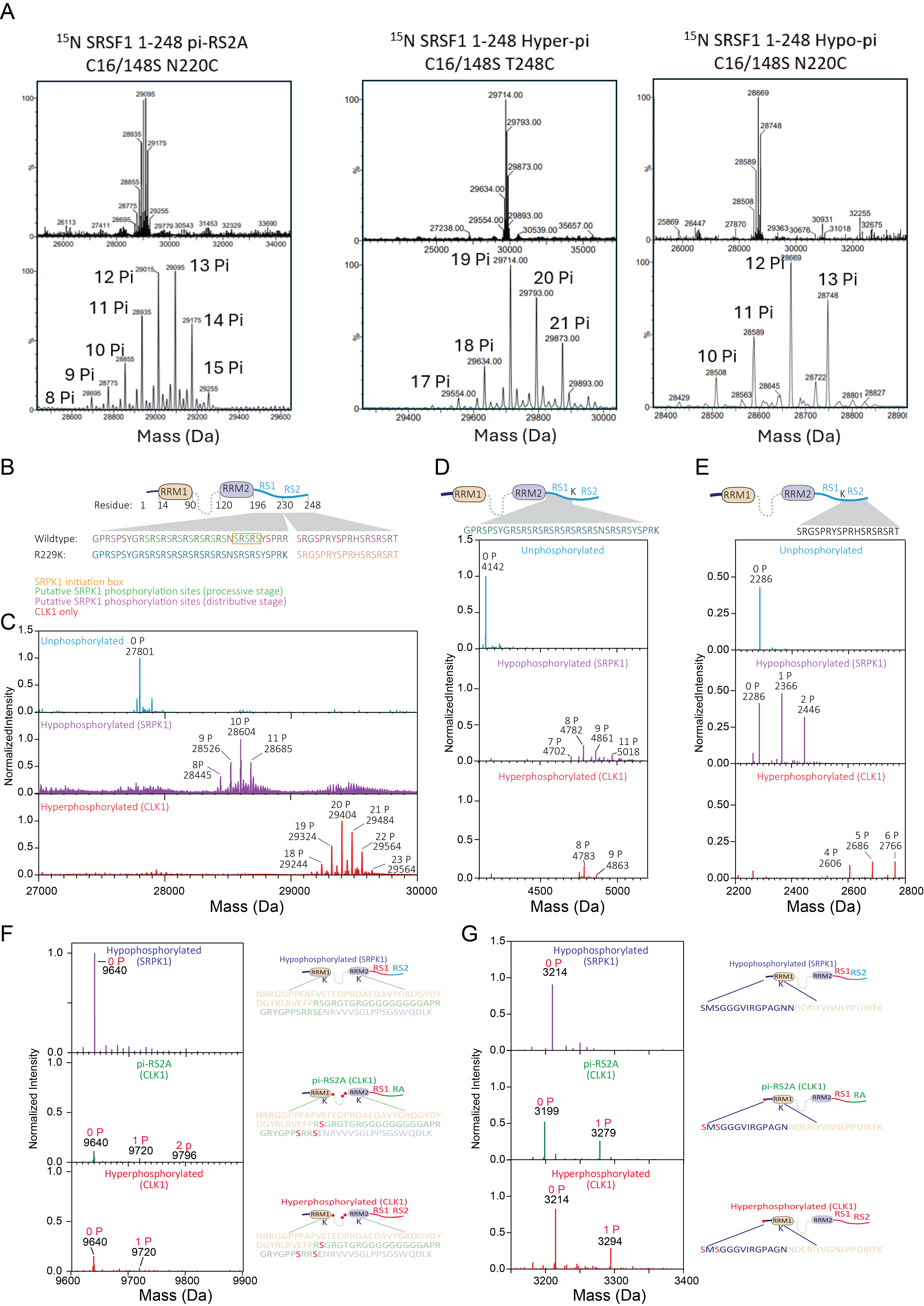

Figure S1. Confirmation of the phosphorylation states for the constructs used in this study. (A) MS of NMR samples for SRSF1 in the un-, hypo-, and hyper-phosphorylated states. Note that the samples were labeled by ^15^N. (B) Design of the R229K construct. R229K is at the boundary of RS1 and RS2 and introduces a cleavage site for Lys-C. SRPK1 binds to an initiation box on the C-terminal end of RS1. Additional phosphates are added in a distributive stage to serines adjacent to arginine residues across both RS1 and RS2. CLK1 has been demonstrated to phosphorylate serines throughout the RS domain. (C) Intact MS of SRSF1 R229K in the unphosphorylated, hypophosphorylated, and hyperphosphorylated states. (D) RS1 peptide and (E) RS2 peptide released by 10-min Lys-C cleavage for the unphosphorylated, hypophosphorylated, and hyperphosphorylated states. (F) The fragment encompassing the RRM1/RRM2 linker obtained after a 12-hour digestion with Lys-C for various SRSF1 constructs placed adjacent to their expected cleavage products. The linker is phosphorylated by CLK1 but not SRPK1. Cleavage product containing the linker residues displays 80 Da shifts upon phosphorylation by CLK1 but not SRPK1. Possible phosphorylation sites are highlighted in red. (G) The N-terminus of the protein is phosphorylated by CLK1 but not by SRPK1. Cleavage product of the most N-terminal fragment of the protein. A species with one phosphate added is present for both CLK1-phosphorylated constructs. Possible phosphorylation sites are highlighted in red.

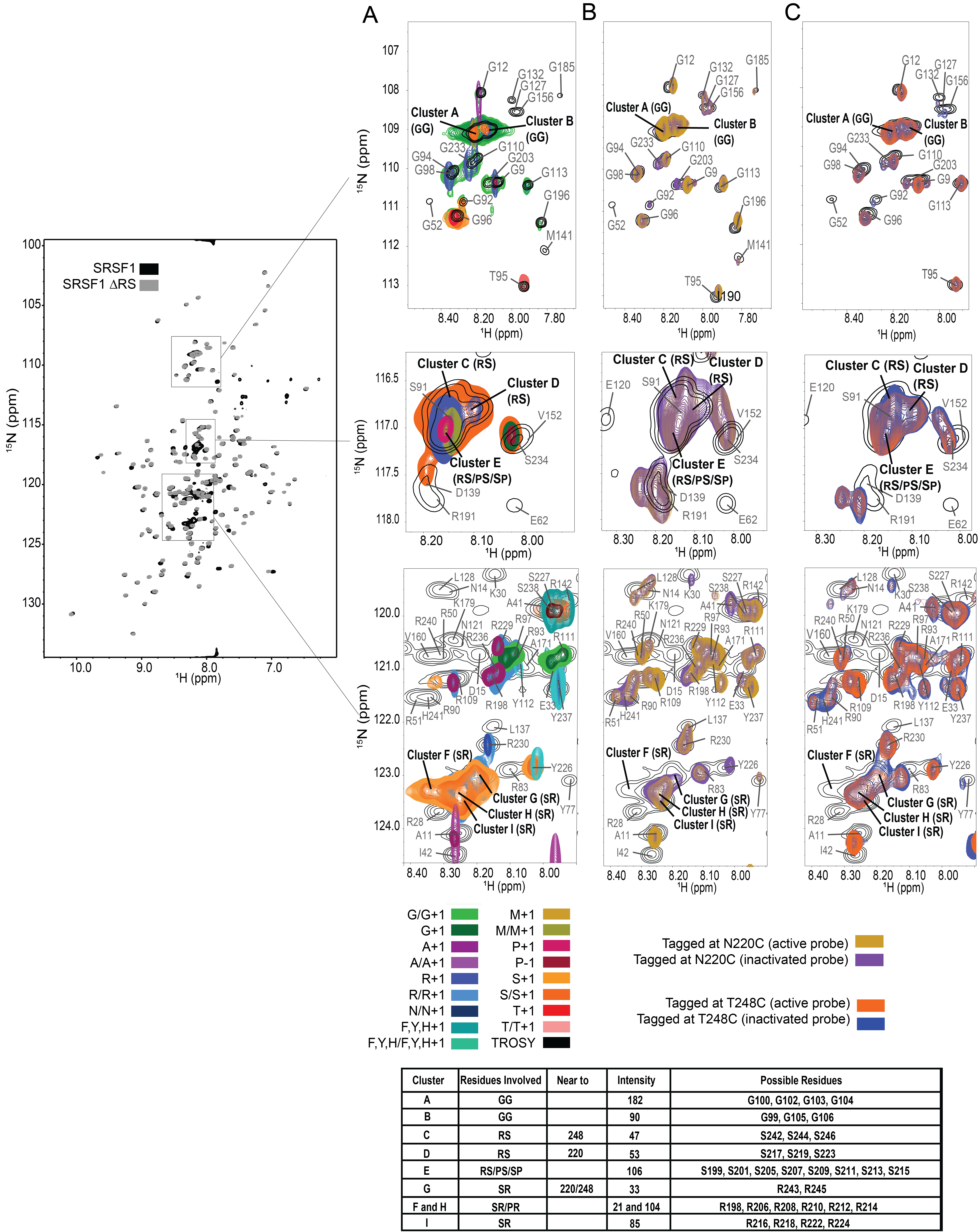

Figure S2. Grouping of unstructured regions of the unphosphorylated protein spectrum based on (A) MUSIC (MUltiplicity Selective In-phase Coherence transfer) as well as HSQC-based PRE spectra with the probe placed at either (B) N220C or (C) T248C.

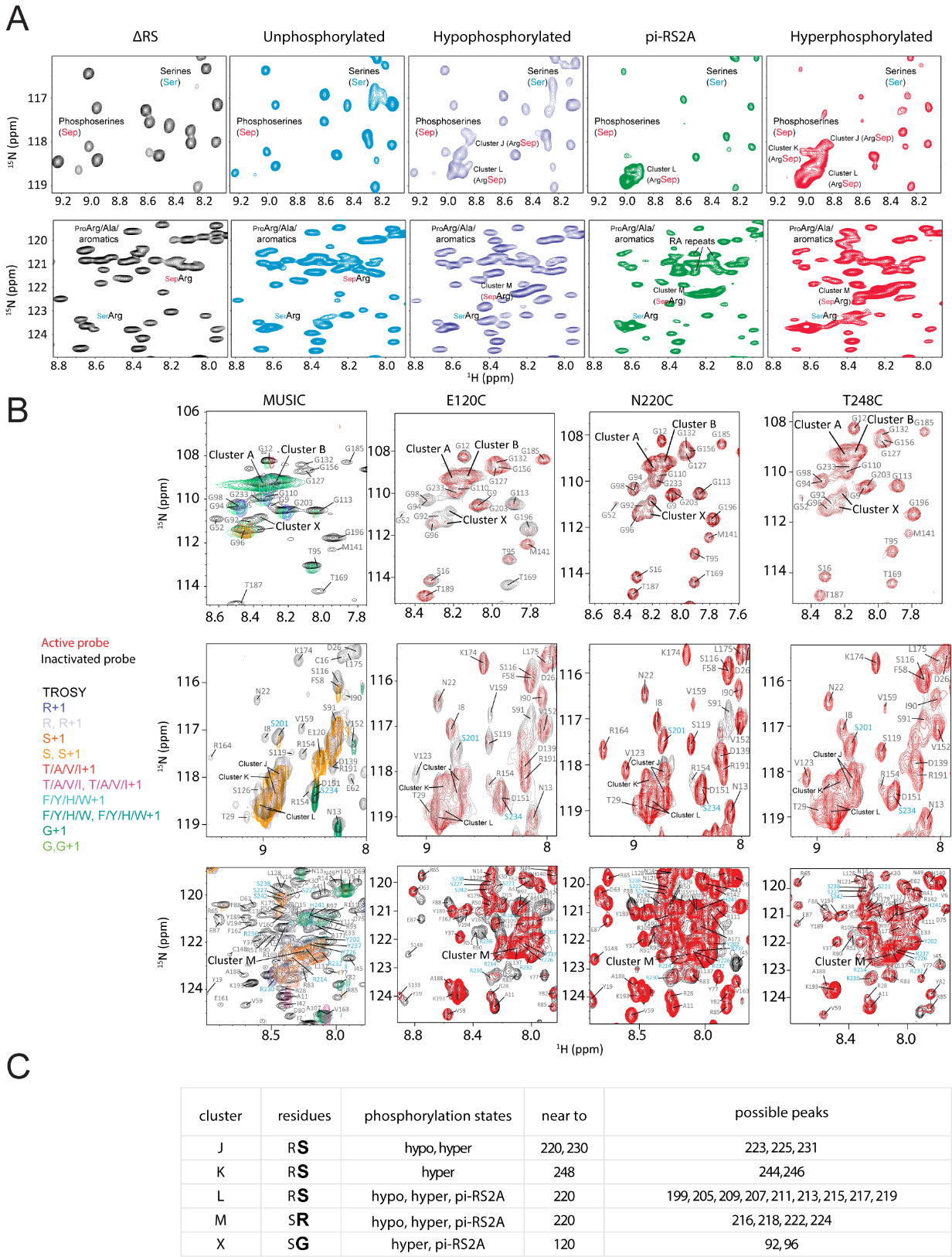

Figure S3. Grouping of unstructured regions of the hyperphosphorylated protein spectrum based on MUSIC (MUltiplicity Selective In-phase Coherence transfer) as well as TROSY-based PRE spectra with the probe placed at either E120C, N220C or T248C. A) Unstructured regions of the spectra of various phosphorylation states. B) Loose assignment of unstructured regions. Peak clusters shown in black. Cyan used to indicate single residues of the RS tail. C) Use of MUSIC and PRE data to group peaks into clusters.

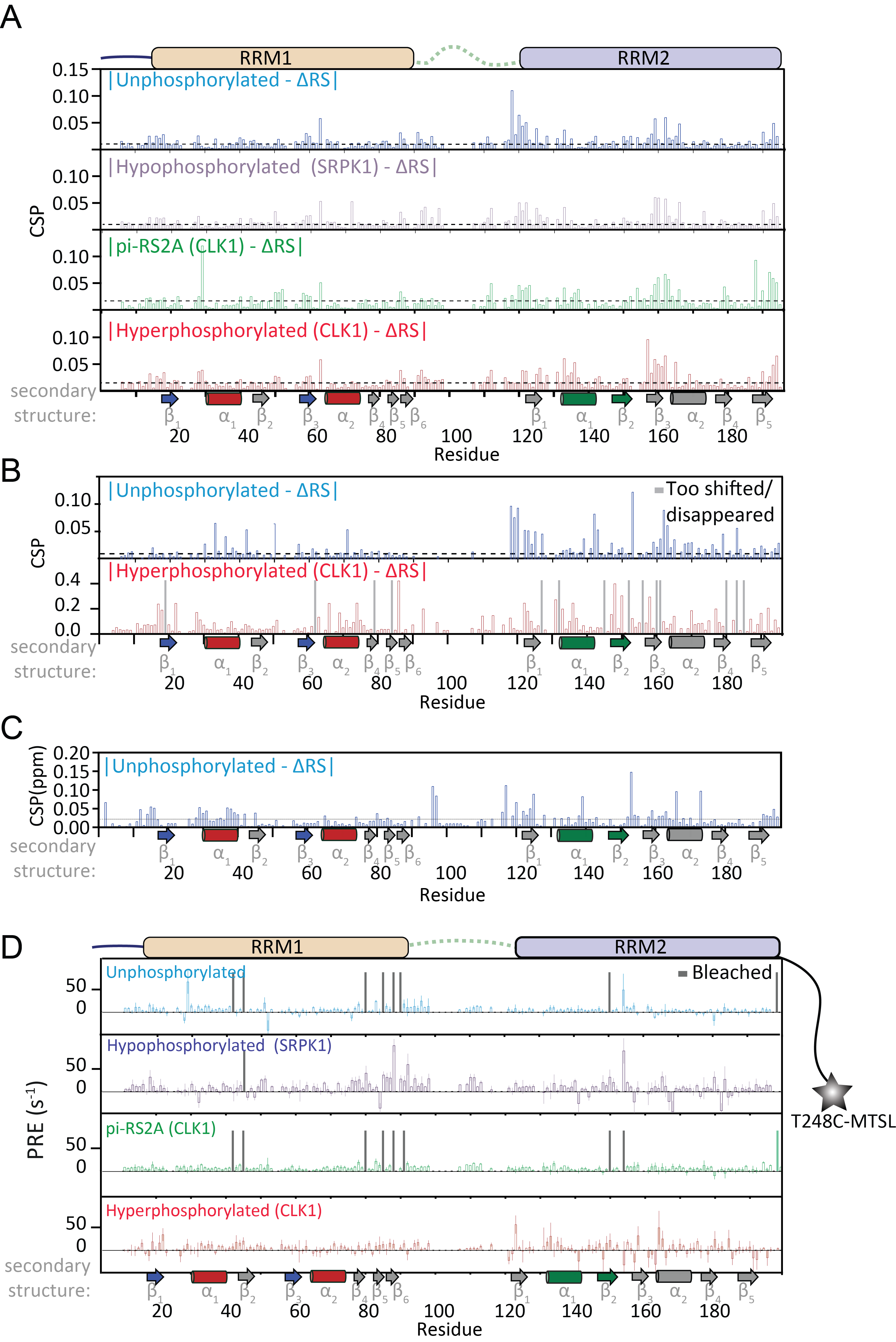

Figure S4. Chemical Shift perturbations and PRE were used to obtain information about interactions with various portions of the RS tail. (A) Chemical shift perturbations in peptide buffer of 100 mM ER4, 400 mM Arg/Glu pH 6.5. (B) Chemical shift perturbations in 200 mM Arg/Glu pH 6.5, (C) Chemical shift perturbations in 46 mM KPO_4_ pH 6.3, 91 mM KCl 50 mM Arg/Glu, 0.91 mM DTT (D) PRE with a paramagnetic probe placed on the very C-terminal end of the protein. Performed in peptide buffer (100 mM ER4, 400 mM Arg/Glu pH 6.5).

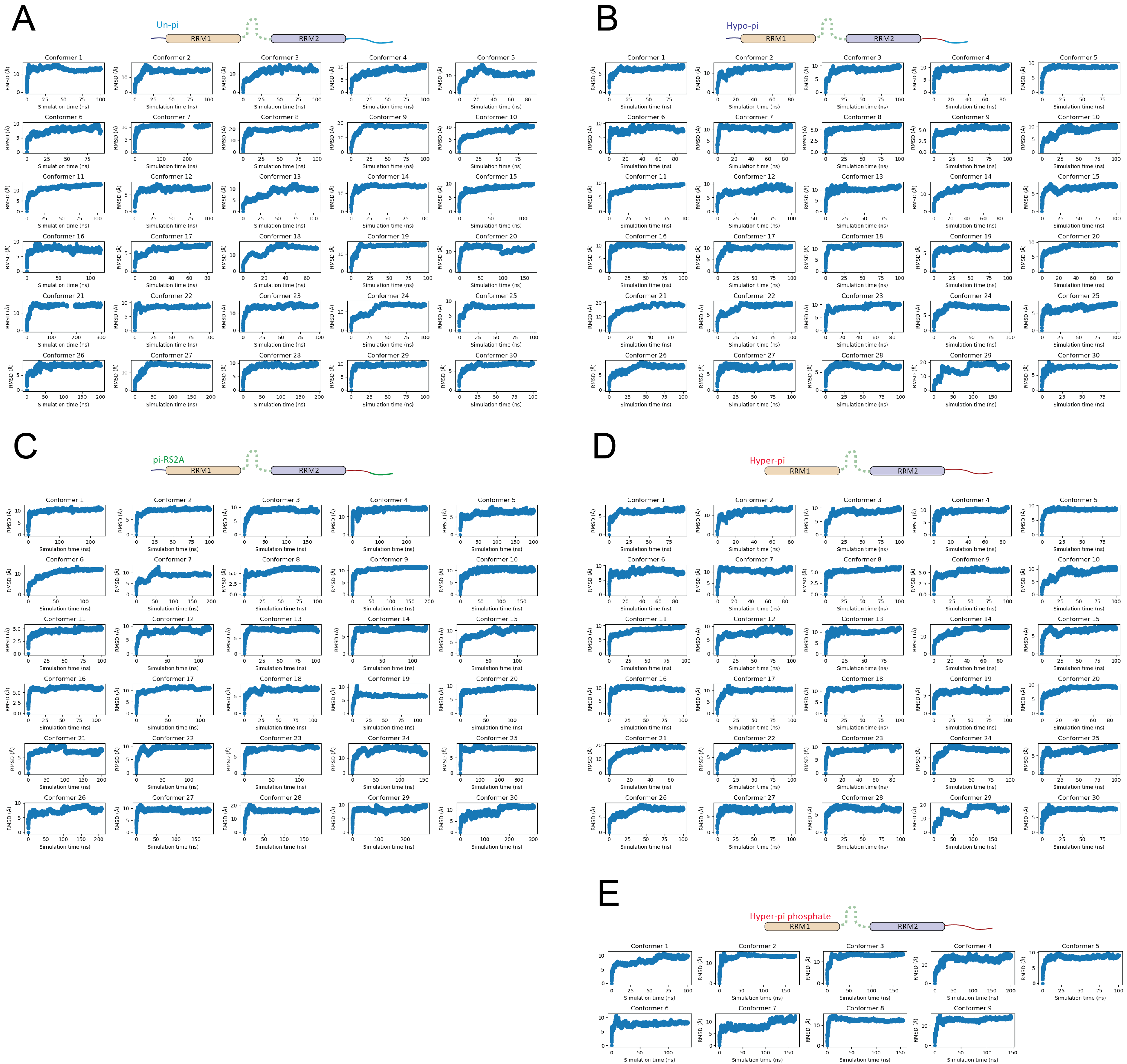

Figure S5. RMSD values between starting coordinates and coordinates of all atoms of the protein over time in the MD simulations for the (A) Unphosphorylated, (B) Hypophosphorylated, (C) pi-RS2A, (D) Hyperphosphorylated constructs refined against PRE in a peptide buffer and (E) Hyperphosphorylated SRSF1 refined against relative PRE obtained in a phosphate buffer.
